# Trafficking of the glutamate transporter is impaired in LRRK2-related Parkinson’s disease

**DOI:** 10.1101/2021.08.04.455053

**Authors:** Ludovica Iovino, Veronica Giusti, Francesca Pischedda, Elena Giusto, Nicoletta Plotegher, Antonella Marte, Ilaria Battisti, Angela Di Iacovo, Algerta Marku, Giovanni Piccoli, Rina Bandopadhyay, Carla Perego, Tiziana Bonifacino, Giambattista Bonanno, Cristina Roseti, Elena Bossi, Giorgio Arrigoni, Luigi Bubacco, Elisa Greggio, Sabine Hilfiker, Laura Civiero

## Abstract

The Excitatory Amino Acid Transporter 2 (EAAT2) accounts for 80 % of brain glutamate clearance and is mainly expressed in astrocytic perisynaptic processes. EAAT2 function is finely regulated by endocytic events, recycling to the plasma membrane and degradation. Noteworthy, deficits in EAAT2 have been associated with neuronal excitotoxicity and neurodegeneration. In this study, we show that EAAT2 trafficking is impaired by the leucine-rich repeat kinase 2 (LRRK2) pathogenic variant G2019S, a common cause of late-onset familial Parkinson’s disease (PD). In LRRK2 G2019S human brains and experimental animal models, EAAT2 protein levels are significantly decreased, which is associated with elevated gliosis. The decreased expression of the transporter correlates with its reduced functionality in mouse LRRK2 G2019S purified astrocytic terminals and in *Xenopus laevis* oocytes expressing human LRRK2 G2019S. In LRRK2 G2019S knockin mouse brain, the correct surface localization of the endogenous transporter is impaired, resulting in its interaction with a plethora of endo-vesicular proteins. Mechanistically, we report that pathogenic LRRK2 kinase activity delays the recycling of the transporter to the plasma membrane *via* Rabs inactivation, causing its intracellular relocalization and degradation. Taken together, our results demonstrate that pathogenic LRRK2 interferes with the physiology of EAAT2, pointing to extracellular glutamate overload as a possible contributor to neurodegeneration in PD.

## Introduction

The concentration of extracellular glutamate in the central nervous system is finely tuned by specific excitatory amino acid transporters (EAATs) [60, 97], which remove the neurotransmitter from the extracellular synaptic milieu and prevent the dramatic consequences of glutamate accumulation [9, 15, 22, 48, 56, 75, 81]. Impaired glutamate uptake entails neuropathological consequences such as alteration of synaptic neurotransmission, neuronal excitotoxicity as well as astro- and microgliosis and neurodegeneration [31, 35]. Among EAATs, EAAT2 (corresponding to glutamate transporter type 1, Glt-1 in rodents) is the predominant glutamate transporter in the adult mammalian brain and accounts for the removal of most of the extracellular glutamate [77, 90]. Approximately 80 to 90 % of EAAT2 is localized on astrocytes, and 5 to 10 % on the axonal terminal of neurons [9, 15, 22, 56, 75, 81].

Regulation of EAAT2 function occurs by protein expression and protein distribution. Indeed, both lateral diffusion and endocytic trafficking ensure a dynamic turnover of the receptor at the synaptic terminals that reflects the shuffling activity of neighboring cells [2, 48, 50, 61]. EAAT2 is constitutively internalized into recycling endosomes *via* a clathrin-dependent pathway, which relies on the reversible ubiquitination of specific lysine residues located in the cytoplasmic C-terminus [27]. The ubiquitin ligase Nedd4-2 has been identified as a mediator of EAAT2 endocytosis [107]. The activation of protein kinase C (PKC) promotes the phosphorylation of Nedd4-2, its association with EAAT2 and the subsequent ubiquitination of the transporter [23]. Importantly, a ubiquitination/de-ubiquitination cycle enables a reversal translocation of EAAT2 from the recycling endosomes back to the plasma membrane. Therefore, a strict regulation of this bidirectional trafficking is crucial to guarantee an appropriate exposure of EAAT2 at the cell surface [27, 51]. Interference with EAAT2 trafficking has been often associated with the re-routing of the transporter to the cellular degradative systems [80, 88, 96, 99].

Parkinson’s disease (PD) is a progressive neurodegenerative disorder clinically characterized by severe motor disability and cognitive impairment [1]. The pathological hallmarks of PD include the selective loss of dopaminergic neurons in the *Substantia Nigra pars compacta* (*SNpc*) that project to the dorsal striatum, the presence of Lewy Bodies in neuronal and glial cells, as well as signs of extended neuroinflammation [32, 83, 84]. Despite decades of research, the mechanisms underlying the selective degeneration of dopaminergic neurons in PD remain unclear [16]. Recent hypotheses support the idea that chronic, subtle dysfunctions proceed years before the clinical symptoms and the dopaminergic death [92]. Perturbation of glutamate homeostasis is one of the earliest events in the pathophysiology of PD and contributes to the exacerbation of later clinical impairments [20, 35, 40, 63, 104]. In addition, deficits in glutamate transporters have been consistently reported upon acute neurotoxin injection in rodents [10, 11, 107] as well as in genetic models of PD [13, 30, 40], and selective ablation of nigral and striatal astrocytic Glt-1 expression induces a parkinsonian-like phenotype in mice [71, 108].

Monogenic mutations with Mendelian transmission have been identified in about 10 % of PD cases [36], and mutations in the leucine-rich repeat kinase 2 (LRRK2) account for up to 40 % of familial PD forms. Specifically, the most common LRRK2 G2019S mutation leads to the expression of a hyperactive form of the LRRK2 kinase and appears in approximately 1 % of apparently sporadic PD cases, with much higher prevalence in specific ethnic groups [93]. Although LRRK2 G2019S animal models do not show clear signs of neurodegeneration, aberrant cortico-striatal glutamatergic neurotransmission has been observed, thus comprising a valuable model system to study the early stages of the disease [8, 46, 52, 53, 95, 98].

Here, we report that the PD-linked LRRK2 G2019S mutation decreases the functionality of EAAT2 in astrocytes by impairing Rab-mediated recycling to the plasma membrane. Overall, our results point to glutamatergic dysregulation as a key pathological event in PD.

## Materials and methods

### Human samples

*Post-mortem* human caudate and putamen were lysed in RIPA buffer (20 mM Tris-HCl pH 7.5, 150 mM NaCl, 1 mM EDTA, 2.5 mM sodium pyrophosphate, 1 mM β-glycerophosphate, 1 mM sodium orthovanadate) containing protease and phosphatase inhibitors (Roche) derived from four LRRK2 G2019S-linked patients, 5 idiopathic PD patients and 5 age-matched controls were obtained from Queen Square Brain Bank (London, UK). *Post-mortem* human brains were collected under human tissue authority license n° 12198. Limited sample demographics are listed in Table 1 and detailed in [49].

**Table 1.**
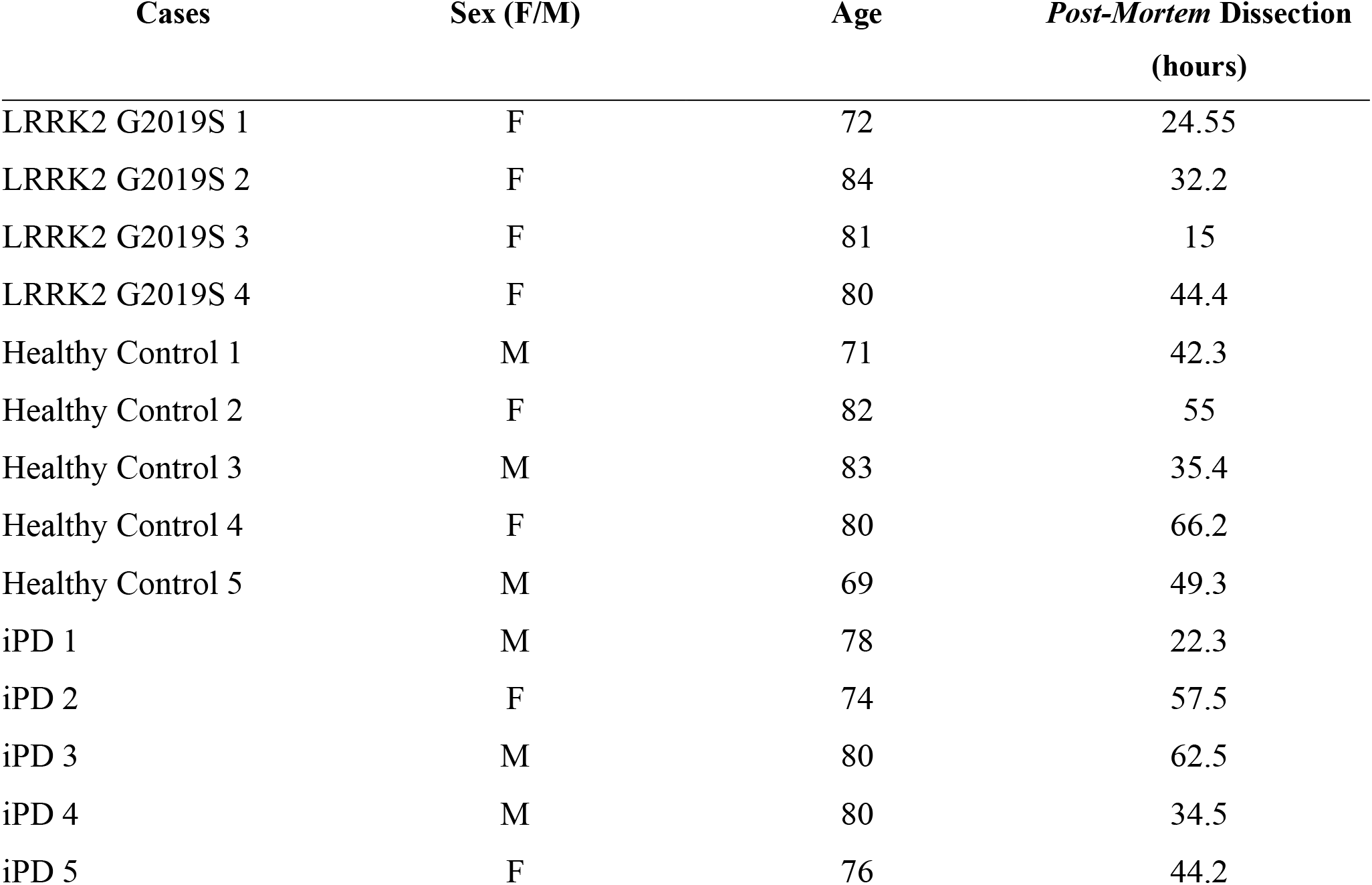
Sample demographics of the human cases used in this study. The table illustrates sex, age and *Post-Mortem* Dissection (expressed in hours) characteristics of LRRK2 G2019S (n=4), Healthy Controls (n=5) and idiopathic PD (iPD; n=5) patients employed in the study.

### Animals

C57Bl/6J Lrrk2 wild-type (WT) and LRRK2 G2019S knock-in mice were used. Lrrk2 G2019S knock-in mice were obtained from Prof. Michele Morari and Novartis Institutes for BioMedical Research, Novartis Pharma AG (Basel, Switzerland) [45]. Housing and handling of mice were done in compliance with national guidelines. All procedures performed in mice were approved by the Ethical Committee of the University of Padova and the Italian Ministry of Health (license 1041/2016, 200/2019 and 105/2019). All procedures performed in *Xenopus laevis* were approved and experiments carried out according to the Ethical Committee of the University of Insubria (license no. 02_15) and the Italian Ministry of Health (1011/2015).

### Immunoblotting

Human caudate and putamen samples as well as dissected striata or primary astrocytes derived from Lrrk2 WT and Lrrk2 G2019S knock-in mice were lysed in RIPA buffer containing 1 % protease inhibitor cocktail (Sigma-Aldrich). Protein concentration was measured using the Pierce® BCA Protein Assay Kit following manufacturer’s instructions (Thermo Scientific). 25 µg protein samples were resolved by electrophoresis on pre-cast 4–20 % tris-glycine polyacrylamide gels (Biorad) and transferred to polyvinylidene difluoride membranes using a semi-dry Biorad transfer machine (Trans-Blot® Turbo TM Transfer System) with the 1X Transfer Buffer (BioRad) at 25 V for 20 min. Membranes were incubated in Tris-buffered saline plus 0.1 % Tween (TBS-T) plus 5 % skimmed milk for 1 h at room temperature (RT), and then incubated overnight with primary antibodies diluted in TBS-T plus 5 % skimmed milk. The following primary antibodies were used: guinea pig anti-glutamate transporter (Glt-1/EAAT2; AB1783, EMD Millipore, 1:500), mouse anti-GAPDH (CSB-MA000195, Cusabio, 1:3000), mouse anti-β-actin (A1978, Sigma-Aldrich, 1:10000), mouse anti-HSP70 (H5147; Sigma Aldrich, 1:5000), mouse anti-Flag-HRP (A8592; Sigma Aldrich, 1:5000), rabbit anti-glial fibrillary acidic protein (GFAP; Z0334, Dako, 1:20000/1:100000), rabbit anti-glutamine synthetase (GS; GTX109121, GENETEX, 1:1000), rabbit anti-Tyrosine Hydroxylase (TH; AB152, EMD Millipore, 1:1000), rabbit anti-Rab8A (ab188574; Abcam, 1:2500), rabbit anti-Rab10 (ab104859; Abcam, 1:1400), rabbit anti-phospho-Rab8A T72 (ab230260; Abcam, 1:1000), rabbit anti-phospho Rab10 T73 (ab230261; Abcam, 1:400), rabbit anti-LRRK2 (MJFF2; Abcam, 1:300), rabbit anti-pS935 LRRK2 (ab133450; Abcam, 1:300) and rabbit anti-pS1292 LRRK2 (ab203181; Abcam, 1:500). Membranes were subsequently rinsed and incubated for 1 h at RT with the appropriate HorseRadish-Peroxidase (HRP)-conjugated secondary antibodies (Invitrogen). The visualization of the signal was conducted using Immobilon® Forte Western HRP Substrate (Millipore) and the VWR® Imager Chemi Premium. Images were acquired and processed with ImageJ software to quantify total intensity of each single band.

### Human caudate and putamen immunofluorescence and immunohistochemistry analysis

Paraffin embedded sections (8 μm) from caudate and putamen were cut using a microtome (Thermo Scientific). To remove paraffin wax, 8 μm-slices were washed in xylene and rehydrated in 100 % ethanol. Slices were treated for 15 min with a quenching solution containing 50 mM NH_4_Cl in phosphate buffer saline (PBS) for immunofluorescence analysis, or 100 % methanol and 3 % H_2_O_2_ for immunohistochemistry studies. Slices were then washed in TBS-T and incubated at 90° C for 15 min with a citrate buffer (10 mM Sodium Citrate and 0.1M citric acid dissolved in distilled water; pH 7.0) to perform epitope retrieval. Subsequently, slices were saturated for 1 h at RT in a blocking solution containing 1 % bovine serum albumin (BSA), 15 % goat serum, 0.25 % gelatin, 0.20 % glycine and 0.5 % Triton. Slices were then incubated overnight at 4 °C with the following primary antibodies diluted in blocking solution: guinea pig anti-EAAT2 (1:200) and/or rabbit anti-GFAP (1:400). The following day, slices were washed in TBS-T and incubated for 1 h at RT with a mixture of the following secondary antibodies diluted 1:200 in blocking solution: anti-guinea pig Alexa Fluor 488 (Life Technologies) and anti-rabbit Alexa Fluor 568 (Life Technologies) fluorophores. Upon washes in TBS-T, slices were incubated for 5 min with Hoechst (1:10000; Invitrogen) to visualize the nuclei and mounted with Mowiol (Calbiochem).

For immunohistochemistry analysis, sections processed with rabbit anti-GFAP were incubated for 1 h at RT using secondary HRP-conjugated anti-rabbit antibody (1:200). Slices were then washed and developed for 2-3 min using 3,3′-diaminobenzidine (DAB) staining kit (Abcam; #ab64238;). Nuclei were counterstained using Haematoxylin (MHS32-Sigma) for 5 min. Finally, sections were dehydrated in graded ethanol (70 %, 90 % and 100 %), transferred in xylene and mounted using a Eukitt Mounting resin (Kirsker Biotech).

### Plasmids

The pCMV-hEAAT2 plasmid encoding the human excitatory amino acid transporter type 2 (EAAT2) was a gift from Susan Amara (Addgene plasmid # 32814; http://n2t.net/addgene:32814; RRID:Addgene_32814). The cDNA was subcloned into pcDNA3 between KpnI and XbaI restriction sites and under the T7 promoter for expression in *Xenopus laevis* oocytes. The pDESTN-SF-TAP LRRK2 plasmids encoding the human WT and G2019S variant were described in [4]. The pCMV6-mGLT-1 (Myc-DDK-tagged) plasmid encoding the mouse glial high affinity glutamate transporter member 2 (Slc1a2), transcript variant 1, was purchased from OriGene Technologies Inc (Cat. MR226166). For the pEGFP-rat Glt1 plasmid, the 99-1820 bp fragment of the Rat Glt-1 plasmid (GenBankTM accession number X67857.1) was subcloned between the EcoRI and Xba restriction sites of pEGFP vector (Clontech). GFP-Rab4, GFP-Rab11 and GFP-Lamp1 plasmids were generated as previously described [26, 73]. RFP-Rab8A WT or Q67L and RFP-Rab10 WT or Q68L plasmids were generated as previously described [42, 47].

### Electrophysiological recordings in *Xenopus laevis* oocytes

pcDNA3_hEAAT2 and pDESTN-SF-TAP-LRRK2 WT and G2019S plasmid vectors were linearized by HindIII and SmaI, respectively. Corresponding cRNAs were transcribed *in vitro* and capped using T7 RNA polymerase. Oocytes were obtained by laparotomy from adult female Xenopus laevis (Envigo). Frogs were anesthetized by immersion in MS222 1 g/L solution in tap water adjusted at final pH 7.5 with bicarbonate. After the treatment with an antiseptic agent (povidone-iodine 0.5 %), the frog abdomen was incised, and the portions of the ovary removed. Oocytes were treated with 1 mg/mL collagenase IA (Sigma Collagenase from Clostridium histolyticum) in calcium-free ND96 (96 mM NaCl, 2 mM KCl, 1 mM MgCl_2_, 5 mM 4-(2-hydroxyethyl)-1-piperazineethanesulfonic acid (HEPES); pH 7.6) for at least 1 h at 18 °C. Healthy and fully grown oocytes were selected and manually separated in NDE solution (ND96 plus 2.5 mM pyruvate, 0.05 mg/mL gentamicin sulfate and 1.8 mM CaCl_2_). After 24 h, healthy looking stage V and VI oocytes were collected and co-injection of cRNA EAAT2 (25 ng) + cRNA LRRK2 WT (25 ng) or cRNA EAAT2 (25 ng) + LRRK2 G2019S (25 ng) was carried out using a manual microinjection system (Drummond Scientific Company). Oocytes were subsequently incubated at 18 °C for 2-3 days in NDE solution [6]. Transport currents (I) were recorded from voltage-clamped oocytes using two microelectrodes filled with 3 M KCl (Oocyte Clamp OC-725C, Warner Instruments). Bath electrodes were connected to the experimental oocyte chamber via agar bridges (3 % agar in 3 M KCl). The external control solution had the following composition: 98 mM NaCl, 1 mM MgCl_2_, 1.8 mM CaCl_2_, 5 mM HEPES, adjusted to pH 7.6 with NaOH. Signals were filtered at 0.1 kHz and sampled at 200 Hz or 0.5 kHz and at 1 kHz. Transport-associated currents were calculated by subtracting the traces in the absence of substrate from those in its presence. To measure the apparent affinity for glutamate (the concentration of glutamate that yields one-half of the maximal transport current), oocytes were exposed to different neurotransmitter concentrations (10 µM, 25 µM, 100 µM, 500 µM, 1 mM). Clampex and Clampfit 10.7 (Molecular Devices) were used to run the experiments, acquire and analyze the data.

### Immunodetection in *Xenopus laevis* oocytes

Injected oocytes were fixed in ice-cold 4 % paraformaldehyde (PFA) in PBS pH 7.5, for 15 min at 4 °C. Oocytes were subsequently washed using ND96 in mild agitation (5 min at RT, three times), included in Polyfreeze tissue freezing medium (Polysciences, Eppelheim) and frozen in liquid nitrogen. Oocyte cryosections (10 μm thickness) were obtained with a cryostat (Leica Biosystems) and preserved at -20 °C. Before use, oocyte slices were washed in PBS for 10 min at RT and incubated in blocking solution (2% BSA (w/v), 0.1 % Tween in PBS) for 45 min. Slices were then incubated overnight at 4 °C with the primary antibody guinea pig anti-EAAT2 (1:200) diluted in blocking solution. The following day, oocyte sections were washed in PBS and incubated with the secondary antibody anti-guinea pig Alexa Fluor 488 fluorophore diluted 1:200 in blocking buffer for 1 h. Sections were washed and mounted using Mowiol. Images were acquired at 8-bit intensity resolution over 1024x1024 pixel on a Leica SP5 confocal microscope using a HC PL FLUOTAR 40x/0.70 oil objective. Using ImageJ, the integrated fluorescence density (IntDen, area x mean fluorescence) of EAAT2 was measured by defining a ROI (8 x 10^-3^ mm^2^ area). All quantifications were applied at the animal pole of the oocytes.

### Free-floating mouse brain slice immunofluorescence

Mice were anesthetized with xylazine (Rompun®) and ketamine (Zoletil®) and transcardially perfused with physiological solution (0.9 % NaCl in PBS) followed by ice-cold 4% PFA dissolved in PBS at pH 7.4. Brains were post-fixed at 4 °C for 18 h in 4 % PFA and then moved in two different solutions of sucrose (20 % and 30 % sucrose in PBS) for 18 h each. Coronal striatal sections (30 μm thickness) were sliced using a vibratome (Campden Instruments Ltd.). Slices were stored at 4 °C in a solution containing 18 % sucrose, 0.01 % NaN_3_ dissolved in PBS. Before staining, murine slices were rinsed in PBS and treated with 50 mM NH_4_Cl dissolved in PBS for 15 min at RT to quench intrinsic autofluorescence. Sections were then washed in PBS followed by permeabilization and saturation for 1 h at RT in a blocking solution containing 2 % BSA, 15 % goat serum, 0.25 % gelatin, 0.2 % glycine and 0.5 % Triton. Slices were then incubated overnight with primary antibodies diluted in blocking solution. The following primary antibodies were used: guinea pig anti-Glt-1 (1:200) and rabbit anti-GFAP (1:400). The following day, slices were washed and incubated for 1 h at RT with the appropriate secondary antibodies diluted in blocking solution: anti-guinea pig Alexa Fluor 488 or 568 fluorophores and anti-rabbit Alexa Fluor 568 or 633 fluorophores. Upon incubation, slices were washed with PBS and nuclei were counterstained with Hoechst 1:10000 and mounted on a glass microscope slide (ThermoFisher) using Mowiol.

Images were acquired at 8-bit intensity resolution over 1024x1024 pixel on a Leica SP5 confocal microscope using a HC PL FLUOTAR 40x/0.70 oil objective. Using ImageJ, the integrated density of GFAP signal upon setting scale and threshold were measured. The number of GFAP^+^ cells was manually counted using the same software.

### Gliosome purification

Glial perisynaptic processes (gliosomes [7, 62, 85]) derived from 4 month old Lrrk2 WT and Lrrk2 G2019S mouse striata were used. Gliosomes were prepared as previously described [65]. Briefly, striata were homogenized in 0.32 M sucrose, buffered at pH 7.4 with Tris-HCl, using a glass-teflon tissue grinder (clearance 0.25 mm – Potter-Elvehjem VWR International). The homogenate was centrifuged (5 min, 1000 g) to remove nuclei and debris, and the supernatant was gently layered on a discontinuous Percoll gradient (2 %, 6 %, 10 %, and 20 % v/v in Tris-buffered 0.32 M sucrose; Sigma-Aldrich). After centrifugation at 33’500 g for 5 min, the layer between 2 % and 6 % Percoll (gliosomal fraction) was collected and washed with a physiological medium having the following composition: 140 mM NaCl, 3 mM KCl, 1.2 mM MgSO_4_, 1.2 mM NaH_2_PO_4_, 5 mM NaHCO_3_, 1.2 mM CaCl_2_, 10 mM HEPES, 10 mM glucose, pH 7.4, and centrifuged at 20000 g for 15 min. The pellet was resuspended in a physiological medium. All the above procedures were conducted at 4 °C. Protein concentration was measured using the Pierce® BCA Protein Assay Kit following manufacturer’s instructions.

### Glutamate uptake assay in gliosomes

Glutamate uptake was measured in striatal gliosomes derived from Lrrk2 WT and Lrrk2 G2019S mice as previously described [58]. Briefly, aliquots of the gliosomal suspensions (500 µL, corresponding to 10-15 µg protein per sample) were incubated for 10 min at 37 °C in the presence of 10 µM 2-amino-5,6,7,8-tetrahydro-4-(4-methoxyphenyl)-7-(naphthalen-1-yl)-5-oxo-4H-chromene-3-carbonitrile (UCPH-101; Abcam 120309, UK), a specific excitatory amino acid transporter 1 (EAAT1/Glast) inhibitor [17]. Then, 20 µL of a solution of [^3^H]D-Aspartate ([^3^H]D-Asp; specific activity: 16.5 Ci/mmol; Perkin Elmer) and non-radioactive D-Aspartate (Sigma Aldrich) was added to each sample to obtain the final concentrations of 0.03, 0.1, 1, 3, 30 and 100 µM. Incubation was continued for further 2 min. Gliosomal samples were exposed to the above [3H]D-Aspartate concentrations for 2 min at 0–4 °C, in the presence of 50 µM of the broad spectrum glutamate transporter blocker DL-threo-beta-benzyloxyaspartate (DL-TBOA; Tocris Bioscience; [82]), to determine the non-specific binding to gliosome membranes. After the 2-min exposure to [^3^H]D-Aspartate, uptake was blocked by rapid vacuum filtration (GF-B filters, Millipore) and filters were washed three times with 5 mL of physiological medium to remove the excess of radioactivity. Radioactivity was determined by liquid scintillation counting. The specific uptake was calculated by subtracting non-specific binding from the total filter radioactivity.

### Primary striatal astrocytes

Mouse primary striatal astrocytes were obtained from postnatal animals between day 1 and day 3 as described in [87]. Brains were removed from their skull and placed in a dish containing cold Dulbecco’s Phosphate Buffered Saline (DPBS, Biowest). Olfactory bulbs and cortices were removed under an optic microscope and striata were transferred to a separate dish containing cold DPBS. After the dissection, Basal Medium Eagle (BME, Biowest), supplemented with 10 % Fetal Bovine Serum (FBS, Corning), 100 U/mL Penicillin + 100 µg/mL Streptomycin (Pen-Strep; Life Technologies), was added to the tissues. Striata were then sifted through a 70-μm cell strainer (Sarstedt) using a syringe plunger. The cell suspension was centrifuged (300 x g, 15 min) and the pellet was washed twice with 25 mL of supplemented medium. Cells were seeded at a density of 5x10^6^ cells/10 mL in cell culture T75 flasks. The culture medium was changed after seven days and again after an additional 3-4 days. When cell confluency reached about 80 %, microglia were detached by shaking the flask (800 rpm) for 2 h at RT. After shaking, the medium containing microglia was replaced with fresh medium. Cells were maintained in BME supplemented with 10 % FBS and Pen-Strep at 37 °C in a controlled 5 % CO_2_ atmosphere. After 14 days, astrocytes were used for the experiments. Each experiment has been performed in triple using at least two independent cell cultures derived from different pups.

### Cell transfection

Astrocytes were seeded at a density of 1x10^5^ cells on 12 mm glass coverslips (VWR) coated with poly-L-lysine. Once 80 % of confluency was reached, cells were transfected using Lipofectamine 2000 (Thermo Scientific) (1:3 DNA/Lipofectamine ratio). For single transfection experiments, 1 µg Myc-DDK-tagged-GLT-1 plasmid was used per each well; for double transfection experiments, 0.5 µg Myc-DDK-tagged-GLT-1 was combined with either 0.5 µg GFP-Rab4, GFP-Rab11 or GFP-Lamp1 plasmids; for triple transfection experiments, 0.3 µg Myc-DDK-tagged-GLT-1 was combined with 0.3 µg GFP-Rab4 and 0.3 µg RFP-Rab8/10 WT or QL plasmids. For TIRFM experiments, astrocytes were seeded at a density of 4x10^5^ cells in 24 mm glass coverslips (VWR) coated with poly-L-lysine. Once at 80 % confluence, cells were transfected with 2.5 µg GFP-GLT-1 plasmid using Lipofectamine 3000 (Invitrogen) (1:2 DNA/Lipofectamine ratio). All experiments were carried out 48 hours after transfection.

### Immunostaining of cultured astrocytes

Cells were fixed using 4 % PFA for 20 min at RT and washed in PBS. Astrocytes were subsequently permeabilized for 20 min in PBS containing 0.1 % Triton and blocked for 1 h at RT in a blocking solution containing 5 % FBS in PBS. For experiments with endogenous Glt-1, astrocytes were incubated with the guinea pig anti-Glt1 (1:200) primary antibody diluted in blocking solution. To stain transfected GFP-Glt-1, astrocytes were incubated with rabbit anti-Glt-1 (1:100) primary antibody [66], whereas transfected Flag-Glt-1 cells were incubated overnight with a mouse anti-Flag® primary antibody (1:200-Sigma Aldrich; #F1804) that recognized the DKK sequence. Co-stainings were performed with a rabbit (1:400) or mouse anti-GFAP (Sigma Aldrich, Cat. SAB5600060-RM246) or a rat anti-Lamp1 (1:300-Abcam; #ab25245) antibody diluted in blocking solution, respectively. The next day, cells were incubated for 1 h at RT with a mixture of anti-guinea pig Alexa Fluor 488 and anti-rabbit Alexa Fluor 568 or 544 secondary antibodies, or with a mixture of anti-mouse Alexa Fluor 568 (1:200) and anti-rat Alexa Fluor 633 in blocking solution. Nuclei were counterstained for 5 min with Hoechst (1:10000), and after washing, coverslips were mounted using Mowiol.

### Organotypic slice culture preparation

Organotypic coronal slices were prepared from 6 up to 8-day-old postnatal pups from C57BL/6J mice as in [78] following the Stoppini’s method [86]. Briefly, pups were anesthetized by hypothermia, the brains were removed from the skull and separated into two hemispheres. Each hemisphere was cut into coronal slices of 400-μm thickness under sterile conditions using a McIlwain tissue chopper. The selected slices were cultured to air-fluid interface-style making use of Millicell culture inserts (30 mm diameter, 0.4 µm; Millipore) at 37°C and 5% CO_2_ for up to 14 days *in vitro*, changing the medium three times a week.

### Biolistic transfection of organotypic slices

Organotypic coronal slices were transfected using a biolistic approach [38]. At 5 DIV, organotypic cultures were treated with 5 μM Cytosine Arabinoside (C1768; Sigma Aldrich) overnight to reduce glial reaction following transfection. The following day, 1.5 mg of 1 μm-gold microparticles (Biorad) were coated with 15 μg GFP-Rab4 plasmid and slices were transfected making use of the DNA transformation apparatus ADNGun (Beambio). After the shot, the medium was replaced, and the slices were cultured for an additional 7 days before visualization.

### Fluorescence microscopy techniques

GFP-Glt-1 transfected astrocytes were imaged by Total internal reflection fluorescence microscopy (TIRFM). TIRFM was achieved with a Carl Zeiss inverted microscope equipped with an Argon laser at 37 °C using a 100×1.45 numerical aperture (NA) oil immersion objective. Green fluorescence was excited using the 488 nm laser line and imaged through a band pass filter (Zeiss) onto a Retiga SRV CCD camera. Images (12 bit, 696x520 pixel) were recorded with the same acquisition parameters (laser power, exposure time, binning) [14]. After setting the threshold, the mean intensity of the GFP signal in three randomly selected regions of interest (ROIs) was measured using ImageJ. Data were expressed as fold changes over GFP-Glt-1 intensity signal measured in Lrrk2 WT animals.

Flag-Glt-1 transfected astrocytes were imaged at 8-bit intensity resolution over 1920x1440 pixel using a Leica 5000B microscope HC PL FLUOTAR 40x/0.75 dry objective. Three ROIs/cell were defined, and the number of Glt-1 positive clusters (diameter > 1 μm) were counted manually using ImageJ.

Co-transfected Flag-Glt-1 astrocytes were imaged at 8-bit intensity resolution over 2048x2048 pixel, using a Leica SP5 confocal microscope and a HC PL FLUOTAR 40x/0.70 oil objective. After setting the threshold, ROIs were identified as the area of Lamp1, Rab11 and Rab4 and visualized in the green channel (GFP) or in the far red for the endogenous staining of Lamp1. Lamp1, Rab11 and Rab4 IntDen were calculated for each ROI. The same ROIs were then transferred to Glt-1 fluorescence channel (Red) and the IntDen was determined as a measure of Glt-1 co-localization. For the characterization of the recycling compartment, the number of GFP-Rab4-positive dots as well as the area of Rab4-positive vesicles was measured using ImageJ.

Transfected organotypic slices were imaged at 8-bit intensity resolution over 2048x2048 pixel, using Zeiss LSM 900 with Airyscan2 and an EC PLAN-Neofluar 40X/1.30 Oil DIC M27 objective.

### Transmission electron microscopy analysis

Primary striatal astrocytes from Lrrk2 WT and Lrrk2 G2019S mice were seeded in 35 mm dishes containing a 14 mm Gridded Coverslip (MatTek, Life Science) and co-transfected with Flag-Glt-1 and GFP-Rab4. After 48 h of transfection, cells were fixed using 4 % PFA dissolved in PBS and GFP-Rab4 was imaged on a Leica SP5 confocal microscope. Subsequently, cells were incubated in 2.5 % glutaraldehyde dissolved in 0.1 M sodium cacodylate buffer for 1 h at 4 °C and post-fixed in 1 % osmium tetroxide plus 1 % potassium ferrocyanide in 0.1 M sodium cacodylate buffer for 1 h at 4 °C. Finally, cells were embedded in the epoxy resin (Epoxy Embedding Medium kit-Sigma Aldrich) and 70 nm-ultrathin sections were obtained using an Ultrotome V (LKB) ultramicrotome, before being processed using a Tecnai G2 transmission electron microscope (TEM) operating at 100 kV. To identify Rab4-positive structures by TEM, confocal and electron images were overlaid using nucleus shape to improve cell relocation. The diameter and area of endosomal-like structures were measured using ImageJ.

### Pharmacological treatments

The following compounds were used: LRRK2 kinase inhibitor rel-3-[6-[(2R,6S)-2,6-Dimethyl-4-morpholinyl]-4-pyrimidyl]-5-[(1-methylcyclopropyl)oxy]-1H-indazole (MLi-2; 200 nM, 90 min application for the experiments on transfected cells; 120 min application for electrophysiological recordings in oocytes, Tocris; Cat. #5756) [18, 43]; dibutyryl cAMP (dbcaMP; 500 μM; 10 days, ChemCruz; #B0218) [109]; PKC activator phorbol 12-myristate 13-acetate (TPA; 400 nM, 20 min, Sigma Aldrich; Cat. #SLBX8889) [109]; PKC inhibitor Go 6976 (10 µM, 1, 10 and 90 min, Selleckchem) [28, 102]; recycling inhibitor monensin (35 µM, 40 min; Sigma Aldrich; Cat. #M5273)[51]; proteasome inhibitor MG132 (20 µM, 8 hours; Sigma Aldrich; Cat. #M5273 [80]); vacuolar H^+^-ATPase inhibitor bafilomycin A1 (Baf A1; 100 nM, 60 min, Selleck.EU; Cat. #S1413;[67]). For the investigation of Glt-1 recycling kinetics, MLi-2 (200 nM) was acutely applied to Lrrk2 G2019S astrocytes 90 min prior to further pharmacological manipulations. For TIRFM experiment and for the treatment of organotypic slices, MLi-2 (200 nM) was acutely dissolved into the medium 90 min prior to immunofluorescence experiments.

### Image preparation and statistical analysis

All images were prepared for illustration using Adobe Illustrator CS6. Statistical analyses were performed in Prism 7 (GraphPad). Data are expressed as median with interquartile range. An outlier test was applied before the statistical analysis. Gaussian distribution was assessed by D’Agostino & Pearson omnibus and Shapiro-Wilk normality tests. Data including three conditions were analyzed by one-way ANOVA test followed by Tukey’s multiple comparisons test (Gaussian distribution) or Kruskal-Wallis test (non-Gaussian distribution) followed by Dunn’s multiple comparisons test. To compare the effects of different drugs on Lrrk2 WT and G2019S astrocytes, we used the Two-way ANOVA test followed by Tukey’s multiple comparisons test. Statistical analysis on data including two independent groups was performed with the Unpaired t-test (Gaussian distribution) or Mann-Withney test (non-Gaussian distribution). A Paired t-test (Gaussian distribution) was applied for the comparison of co-injected EAAT2+ LRRK2 G2019S oocytes before and after MLi-2 application. Levels of significance were defined as *, $, #, p ≤ 0.05; **, $$, ##, p ≤ 0.01; ***, $$$, ### p ≤ 0.001.

## Data availability

Data supporting the findings of the present study are available from the corresponding author, upon reasonable request.

## Supplementary materials and methods

### Quantitative Polymerase Chain Reaction (qPCR)

Total RNA was extracted from mice striata or from primary striatal astrocytes with the Total RNA Purification kit (NORGEN Biotek) and quantified by absorbance in a NanoDrop 2000c UV-Vis spectrophotometer (ThermoFisher Scientific). cDNA was synthesized with the All-in-One Cdna Synthesis SuperMix (Bimake) following manufacturer’s instructions. Gene expression was quantified by qPCR in real-time PCR reactions with Sybr Green technology in a CFX96 Touch Real-Time PCR Detection System (Bio-Rad). 30 ng cDNA were used in iTaq Universal SYBR Green Supermix (Bio-Rad) at the following conditions: stage 1: 95 °C, 5 min; stage 2: 39 x (95 °C, 15 s; 60 °C, 30 s).

The primers were as follows: mSLC1A2 fw: GGTGGAAAGCCGGGACGTGGATTA; mSLC1A2 rev: GCTTGGGCATATTGTTGGCACCCT; SLC1A3 fw: ATCCGGGAGGAGATGGTGCCCGT; SLC1A3 rev: AGGATGCCCAGAGGCGCATACCACA; ACTIN fw: TACCACCATGTACCCAGGCATT; ACTIN rev: ACTCATCGTACTCCTGCTTGCTGA; mGAPDH fw: GAGAGTGTTTCCTCGTCCCG; mGAPDH rev: ACTGTGCCGTTGAATTTGCC; TRANSFERRIN RECEPTOR fw TATAAGCTTTGGGTGGGAGGCA; TRANSFERRIN RECEPTOR rev AGCAAGGCTAAACCGGGTGTATGA;

Primers for Slc1a2, Slc1a3, β-Actin and Transferrin were purchased from Sigma Aldrich while primers for GAPDH were purchased from Metabion International AG. The quantification of the gene relative expression was carried out by the Delta-Delta Ct method [44] by normalizing to the reference genes GAPDH, β-Actin and Transferrin receptor. A dissociation curve was built in the 60-95 °C range to confirm the specificity of the amplification product.

### Liquid-chromatography mass spectrometry (LC-MS) analysis

Endogenous Glt-1 was immunoprecipitated from homogenized WT and Lrrk2 G2019S mouse striata using 5 µg of a rabbit anti-Glt-1 antibody (Abcam; ab205248) in RIPA buffer (1.5 mg of total protein each). Magnetic beads (Protein A/G Magnetic Beads, BioTool) (50 µL) were used to pre-clear and precipitate the transporter. Beads were then washed three times using 1 ml of RIPA buffer and proteins loaded on pre-cast 4–20 % tris-glycine polyacrylamide gels (Biorad). Gel bands were excised, cut into small pieces and destained with a solution of 60 % NH_4_HCO_3_ 200 mM/40 % acetonitrile (ACN) at 37 °C. Disulfide bridges were reduced with 2 mM Tris (2-carboxyethyl)phosphine hydrochloride (TCEP) in 50 mM NH_4_HCO_3_ at 56 °C for 1 h, and cysteine residues were alkylated with 4 mM methyl methanethiosulfonate (MMTS) with 50 mM NH_4_HCO_3_ in the dark at room temperature for 45 min. Gel samples were washed twice with 50 mM NH_4_HCO_3_ and ACN alternatively and vacuum-dried. Samples were incubated with 12.5 ng/µL trypsin (Sequencing Grade Modified Trypsin, Promega) in 50 mM (NH_4_)_2_HCO_3_ and protein digestion was carried out overnight at 37 °C. Peptides were extracted with three changes of 50 % ACN/ 0.1 % formic acid (FA). Samples were vacuum-dried and stored at −20 °C. Samples were analyzed using a LTQ Orbitrap XL mass spectrometer (Thermo Fisher Scientific) coupled to a HPLC Ultimate 3000 (Dionex - Thermo Fisher Scientific) through a nanospray source (NSI), as described previously [3]. Samples were resuspended in 30 µL of 3 % ACN/0.1 % FA and each sample was acquired twice. Raw data files were analyzed with MaxQuant [12] software (v. 1.5.1.2) interfaced with Andromeda search engine. Protein search was performed against the mouse section of the UniProt database (*Mus musculus*, version 2020-09-30, 55494 entries). Enzyme specificity was set to trypsin with up to one missed cleavage allowed, while methylthio-cysteine and oxidized methionine were set as fixed and variable modifications, respectively. A minimum of two peptides was required for protein identification, and a false discovery rate (FDR) ≤ 0.01, both at the peptide and protein level, was used to filter the results. To estimate the relative protein abundance across samples, the intensity values calculated by the software were used. The datasets were compared to highlight significant differences in the protein abundances: common and unique Glt-1 binders between the Lrrk2 WT and G2019S genetic environments were identified performing a Z-test on the log-transformed values of the IP/CTRL ratios and taking into account only proteins with p < 0.05 and Fold Changes (FC) ≥ 3.5. Gene Ontology (GO) terms were employed to identify discrete functional enrichments in order to extract information about novel networks of interactors in relationship to the different subcellular localization of the transporter. G:profiler (https://biit.cs.ut.ee/gprofiler/gost) was used for enrichment analysis. Glt-1-related protein networks in Lrrk2 G2019S pathogenic background were ranked based on the fold-change affinity over the control (≥3.5).

## Results

### Severe deficits in EAAT2 levels and increased gliosis in post-mortem caudate and putamen of LRRK2 G2019S PD patients

To investigate the role of EAAT2 in the pathophysiology of LRRK2-PD, we first examined the presence of the glutamate transporter protein in the caudate and putamen of LRRK2 G2019S PD patients and age-matched controls by western blot (Fig. 1A). Consistent with previous reports, the EAAT2 antibody recognized multiple bands corresponding to the monomeric (60 KDa), and multimeric, SDS-resistant (180 KDa) conformations [24, 55, 111]. An additional 250 KDa band was detected which is compatible with EAAT2 multimers [25, 91] (Fig. 1A). We determined the total amount of the transporter by combining the densitometry of the three bands (Fig. 1B), and we also quantified the three bands separately (Supplementary Fig.1 A-C). In all cases, band intensity was normalized to the housekeeping protein GAPDH. We observed that EAAT2 protein was nearly absent in the caudate and putamen of LRRK2 G2019S PD patients as compared to healthy controls (Fig. 1B; LRRK2 G2019S PD patient cases vs age-matched controls; p=0.01), and this difference affected both the multimeric and monomeric fractions (Supplementary Fig. 1A-C; LRRK2 G2019S PD patients vs age-matched controls: A, p=0.004; B, p=0.006; C, p=0.012). To assess whether EAAT2 downregulation was specifically linked to LRRK2-related pathology, we compared the expression of the transporter in the caudate and putamen of age-matched controls, LRRK2 G2019S and idiopathic Parkinson’s disease (iPD) patients (Supplementary Fig. 1D). The total EAAT2 expression showed a modest but non-significant decrease in the iPD group (Supplementary Fig. 1E-LRRK2 G2019S PD cases vs age-matched controls; p=0.001; LRRK2 G2019S PD cases vs iPD cases; p=0.07; age-matched controls vs iPD cases; p>0.99). The quantification of the monomeric (60 KDa) and multimeric (180 and 250 KDa) EAAT2 bands also confirmed a subtle but non-significant EAAT2 deficit in the iPD samples (Supplementary Fig. 1F-H; iPD cases versus age-matched controls; p>0.05).

**Figure 1.**
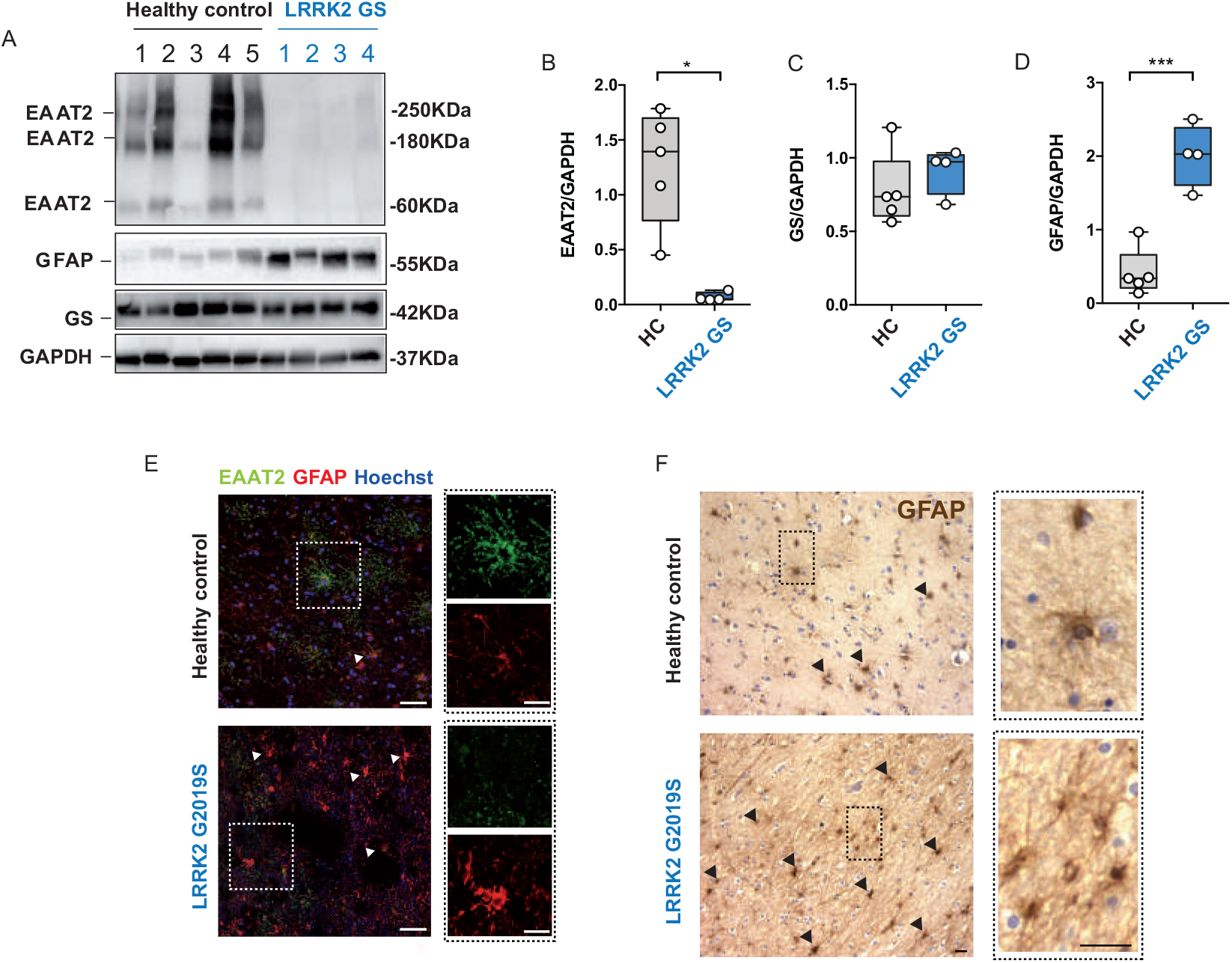
EAAT2 levels are decreased in human LRRK2 G2019S brains. A) Western blot analysis of human LRRK2 G2019S caudate and putamen lysates and healthy controls using anti-EAAT2, anti-GS and anti-GFAP antibodies; B-D) Relative quantification of band intensity was performed using ImageJ and normalized to the housekeeping protein GAPDH (n=5 age-matched control samples and n=4 LRRK2 G2019S caudate and putamen samples); E) Representative double-labeling images for EAAT2 (green) and GFAP (red) in LRRK2 G2019S human caudate and putamen and age-matched control; scale bar 50 μm, insets 20 μm; F) Representative images of DAB-immunostaining for GFAP in LRRK2 G2019S human caudate and putamen and age-matched control; scale bar 25 μm, insets 10 μm; arrowheads in E) and F) indicate the presence of GFAP positive cells. Statistical analysis in B was performed using Mann-Whitney test and in C-D using Unpaired T-test.

EAAT2 is highly expressed by astrocytes [22]. To determine whether the decreased expression of EAAT2 was due to a lower number of astrocytes in LRRK2 G2019S PD patients, we assessed the amount of the cytosolic marker glutamine synthetase (GS), since its expression is restricted to astrocytes and does not depend on their activation status (Fig. 1A). GS expression levels in the caudate and putamen were comparable between LRRK2 G2019S PD patients and age-matched controls (Fig. 1C; LRRK2 G2019S PD patients vs controls, p=0.37). However, expression levels of GFAP (a marker for astrogliosis) were significantly increased in the caudate and putamen of LRRK2 G2019S PD patients as shown in Fig. 1D-F (D, LRRK2 G2019S PD patients vs age-matched controls; p=0.0003). Moreover, immunohistochemical detection of GFAP-positive cells (Fig. 1E-F, arrows) and the quantification of both GFAP IntDen and GFAP^+^ cells (Supplementary Figure 1 I,J) showed that astrocytes in caudate and putamen of LRRK2 G2019S PD patients were abundant and adopted the typical morphology of hyperactive cells, with enlarged and densely stained cytoplasm (Fig. 1E-F, insets). Taken together, these data show a significant decrease in the expression of the glutamate transporter EAAT2 in the caudate and putamen of LRRK2 G2019S PD patients, and the reduction of EAAT2 levels is associated with increased astrogliosis.

### Glutamate transporter expression and functionality are perturbed in the striatum of young LRRK2 G2019S mice

To corroborate the results obtained from a small sampling of LRRK2-PD patients in a defined mouse model, we compared the expression of Glt-1, the mouse equivalent of EAAT2, in the striatum of Lrrk2 WT and Lrrk2 G2019S knock-in mice. Moreover, to inquire whether deficits in glutamate transporter expression occur before dopaminergic degeneration, we chose 4-month old animals since previous studies have reported alterations in glutamatergic neurotransmission in 4 months old mice which correlate with onset of altered motor activity [46, 98]. As previously reported, we confirmed the presence of two bands, corresponding to Glt-1 monomers (60 KDa) and Glt-1 multimers (180 KDa), respectively (Fig. 2A) [24, 55, 111]. Lrrk2 G2019S mice presented a decrease in the total amount of Glt-1 as compared to Lrrk2 WT controls (Fig. 2B, Lrrk2 WT vs G2019S; p=0.008). When analyzing the two bands separately, the lower Glt-1 band also showed a significant decrease in the Lrrk2 G2019S mice, whilst levels of the higher band did not reach statistical significance (Supplementary Fig. 1K, Lrrk2 WT vs Lrrk2 G2019S; p=0.0175; Supplementary Fig. 1L, Lrrk2 WT vs Lrrk2 G2019S; p=0.07).

**Figure 2.**
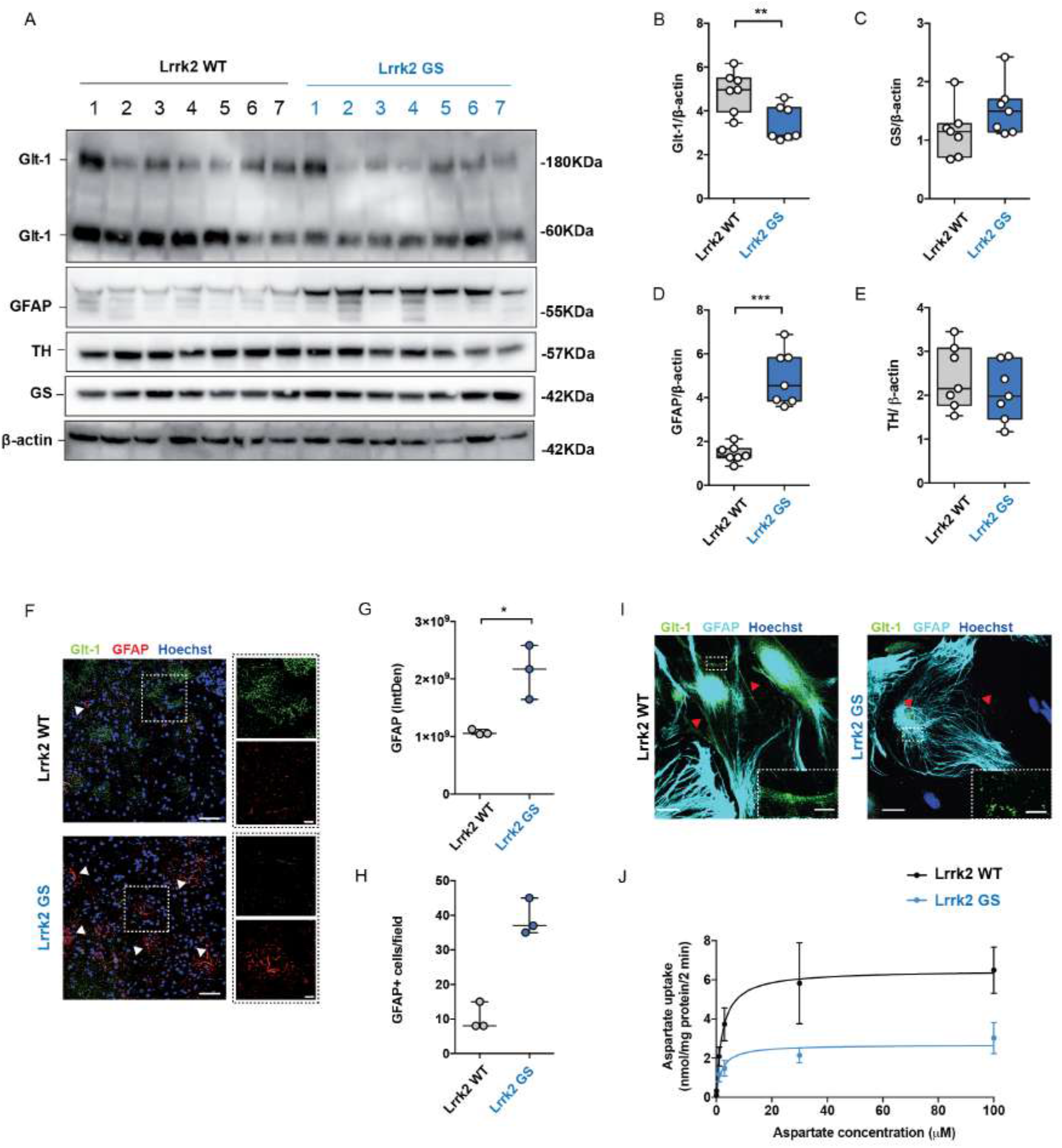
Glutamate transporter is downregulated in the striatum of Lrrk2 G2019S mice. A) Western blot analysis of Lrrk2 WT and Lrrk2 G2019S striatal lysates using anti-Glt-1, anti-GS, anti-GFAP and anti-TH antibodies; B-E) Relative quantification of band intensity was performed using ImageJ and normalized to β-actin (n=7 striatal samples for both Lrrk2 WT and Lrrk2 G2019S 4-month old mice); F) Representative confocal double-labeling images for Glt-1 (green) and GFAP (red) in Lrrk2 WT and G2019S striatal slices; scale bar 50 μm, insets 20 μm; G) Quantification of GFAP IntDen and GFAP^+^ cells (H) in the dorsal striatum of Lrrk2 WT and G2019S mice. Three different fields per animal were collected, n=3 animals for each genotype; I) Representative confocal images of the cellular distribution (red arrowheads) of the endogenous Glt-1 (green) in primary striatal astrocytes stained for GFAP (cyan) derived from Lrrk2 WT and Lrrk2 G2019S mice treated with dbcAMP (500 μM) for ten days; scale bars 20 μm, inserts 5 μm; J) Striatal gliosomes from Lrrk2 WT and Lrrk2 G2019S 4-month old mice were exposed for 2 min at 37 °C to increasing concentration of [^3^H]D-Asp (0.03, 0.1, 1, 3, 30 and 100 µM) in the presence of 10 μM UCPH to exclude [^3^H]D-Asp uptake by Glast. The specific [^3^H]D-Asp uptake is expressed as nmol/mg protein/2 min; the kinetic parameters V_max_ and K_m_ were obtained by fitting data with the Michaelis-Menten equation (n=4 independent experiments for each group). Statistical analysis in B-E and J was performed using Unpaired T-test; statistical analysis in G-H was performed using Mann-Whitney test.

We next performed qPCR analyses to explore whether the decrease in Glt-1 protein levels was attributed to a decrease of its mRNA levels in the striatum. Importantly, no significant changes in mRNA levels between genotypes were detected for the two main glutamate transporters Glt-1 (Supplementary Fig.1M, Lrrk2 WT vs Lrrk2 G2019S; p=0.99) or the glutamate/aspartate transporter (Glast), which is the second most important astrocytic glutamate transporter (Supplementary Fig.1M, Lrrk2 WT vs Lrrk2 G2019S; p=0.6575). Similar to what we observed in human samples, GS protein levels were comparable between the genotypes (Fig. 2C, Lrrk2 WT vs Lrrk2 G2019S, p=0.1411), while there was an increase in the levels of GFAP (Fig. 2D, Lrrk2 WT vs Lrrk2 G2019S; p<0.0001), suggesting enhanced gliosis in young Lrrk2 G2019S mice. Immunofluorescence imaging confirmed a reduction of Glt-1 staining in the striatum of Lrrk2 G2019S mice compared to controls (Fig. 2F). Quantifications of the integrated fluorescence intensity (IntDen) of GFAP (Fig. 2G; Lrrk2 WT vs Lrrk2 G2019S, p=0.0001) and of the number of GFAP-positive astrocytes (Fig. 2H; Lrrk2 WT vs Lrrk2 G2019S, p=0.1) further indicate that decreased Glt-1 levels correlate with an intense astrogliosis in LRRK2 G2019S mice. According with previous reports [39, 69, 106], Tyrosine Hydroxylase (TH) levels as quantified by western blot from the same mice revealed no signs of dopaminergic cell loss, indicating that the decrease in Glt-1 levels seems to precede neurodegenerative events (Fig. 2E, Lrrk2 WT vs Lrrk2 G2019S, p=0.39).

To dissect the LRRK2-mediated effects on glial Glt-1 levels, we isolated primary striatal astrocytes from Lrrk2 WT and Lrrk2 G2019S mice (Fig. 2I). Astrocytes were treated with dbcAMP to stimulate endogenous Glt-1 expression [89] (Fig. 2I). Consistent with published data [79], qPCR analysis showed a more than 10-fold increase in Glt-1 mRNA levels upon dbcAMP treatment (Supplementary Fig. 1N). Immunocytochemistry revealed that Glt-1 was diffusely expressed in Lrrk2 WT astrocytes, while Lrrk2 G2019S astrocytes displayed a less pronounced staining, almost restricted to discrete dots (Fig. 2I, arrowheads).

To evaluate whether the Glt-1 deficits and impaired Glt-1 localization may functionally affect glutamate re-uptake in astrocytes, we purified striatal mouse gliosomes from Lrrk2 WT and Lrrk2 G2019S mice. Gliosomes constitute pure subcellular *ex-vivo* preparations that resemble most of the molecular and functional features of astrocytes *in vivo* [65, 85]. [^3^H]D-Asp, a non-metabolizable analogue of glutamate used to mimic the endogenous neurotransmitter [68], was applied to assess Glt-1 activity, and experiments were conducted in the presence of 10 μM UCPH-101 to exclude [^3^H]D-Asp uptake by Glast. To calculate the K_m_ and the V_max_ of [^3^H]D-Asp uptake, radioactivity was determined in gliosomes in the presence of different concentrations of [^3^H]D-Asp. As shown in Fig. 2J, Lrrk2 G2019S-derived gliosomes displayed a lower V_max_ of aspartate uptake as compared to age-matched Lrrk2 WT mice (Fig. 2J, V_max_ [Lrrk2 WT]: 6.7±1.37 nmol/mg/2 min, V_max_ [Lrrk2 G2019S]: 2.73±0.59 nmol/mg/2 min, p= 0.036), while no significant difference was registered in the K_m_ values (Fig. 2J; K_m_ [Lrrk2 WT] 2.56±0.16 mM, K_m_ [Lrrk2 G2019S] 2.05±0.14 mM, p=0.054). Altogether these findings show that, astrocytes from Lrrk2 G2019S mice present a reduced expression of Glt-1, which results in reduced glutamate uptake and is associated with increased gliosis.

### Human pathogenic G2019S LRRK2 mutation impacts on EAAT2 electrophysiological properties

To confirm that the reduced glutamate uptake observed in LRRK2 G2019S gliosomes also applies to the human EAAT2 protein, we used *Xenopus laevis* oocytes, an excellent model system to study transport processes at a cellular level [103]. Human glutamate transporter EAAT2 mRNA was co-injected with human LRRK2 WT or LRRK2 G2019S mRNA into oocytes, and two-electrode voltage-clamp was performed to record the transport current elicited by glutamate through EAAT2 (Experimental design in Fig. 3A). The inward transport associated-current (I_EAAT2_) obtained upon the application of glutamate to oocytes injected with LRRK2 G2019S mRNA was significantly reduced as compared to the one obtained from oocytes injected with LRRK2 WT mRNA (Fig. 3B; EAAT2+LRRK2 WT: mean amplitude 43±3 nA vs EAAT2+LRRK2 G2019S: mean amplitude 25±2 nA; p<0.0001). We also measured the maximal transport current (I_max_) and the apparent affinity for glutamate (aK_m_). We found that the expression of LRRK2 G2019S mRNA significantly decreased the I_max_ but not the aK_m_ of the transporter as compared to the WT counterpart (Fig. 3C; EAAT2+LRRK2 G2019S, aK_m_: 51±27 µM, I_max_: 28±3 nA vs EAAT2+LRRK2 WT, aK_m_: 89±32 µM, I_max_: 52±5 nA; aK_m_, p =0.05; I_max_, p<0.001).

**Figure 3.**
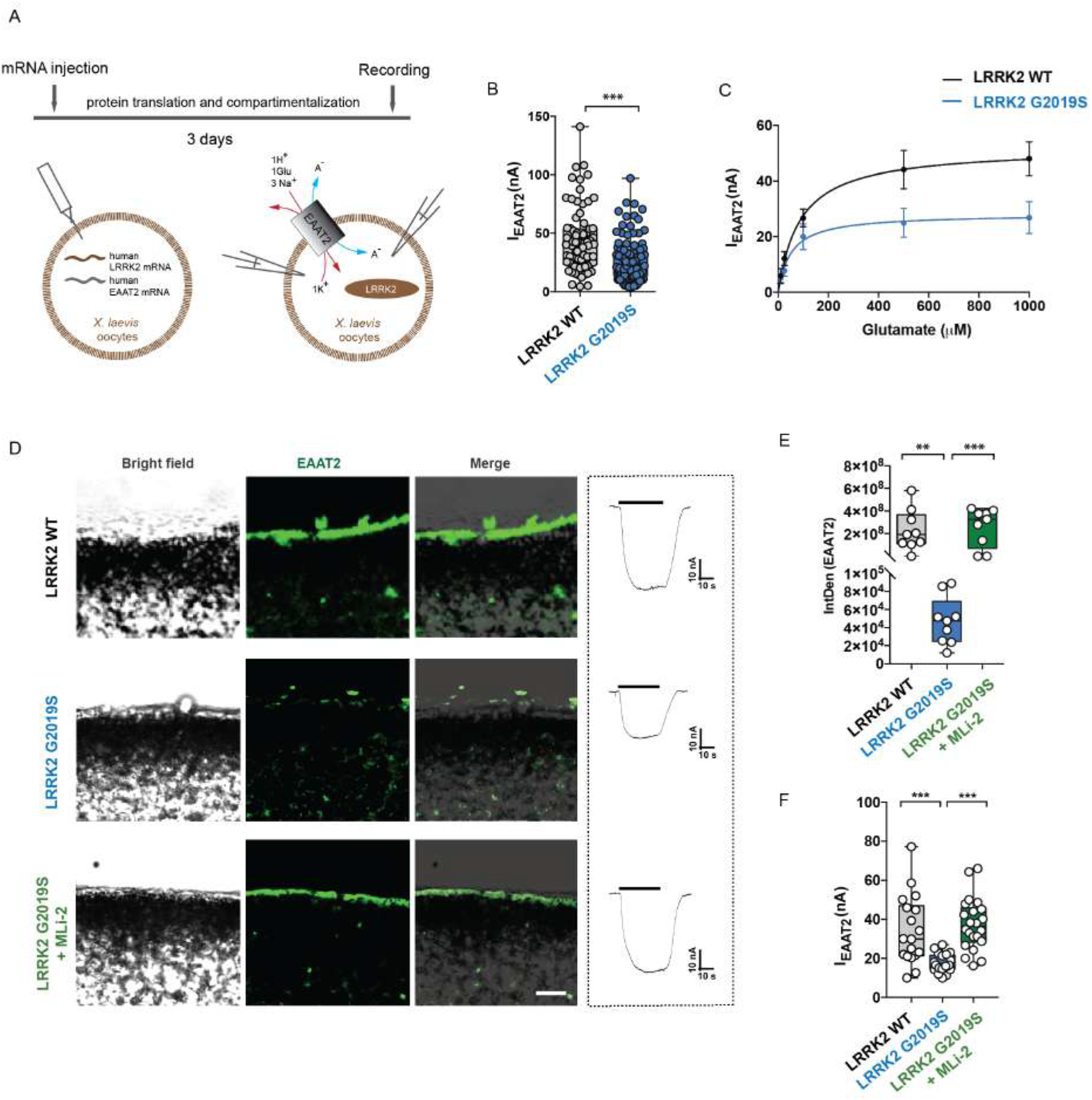
LRRK2 G2019S alters EAAT2 electrophysiological properties. A) Schematic outline of the experimental setup. Oocytes were co-injected with mRNA of EAAT2 and human LRRK2 (WT or G2019S) and glutamate transport associated-currents were recorded using the two-electrode voltage-clamp technique; B) The bar graph represents current amplitudes elicited in oocytes co-expressing EAAT2 and LRRK2 WT (n=79 oocytes, 12 frogs) or G2019S (n=122 oocytes, 12 frogs). Glutamate application was at 1 mM and the holding potential at −60 mV; C) Transport-associated currents in oocytes co-expressing EAAT2 and LRRK2 WT or G2019S as a function of glutamate concentration. The kinetic parameters I_max_ and aK_m_ were obtained by fitting data with the Michaelis-Menten equation (n=6 oocytes; 2 frogs); D) Representative bright field (left column) or fluorescence (middle column, merge on the right column) images of oocyte slices co-expressing EAAT2 (green) and LRRK2 WT or G2019S, with or without 90 min MLi-2 (200 nM) treatment; scale bar 20 μm. Representative traces of the recorded transport current in all the three groups are shown on the right; E) Quantitative analysis of the IntDen of the EAAT2 signal at the oocyte membrane was performed in three different fields for each oocyte; n=3 oocytes for each group; F) The bar graph represents current amplitudes elicited in oocytes co-expressing EAAT2 and LRRK2 WT (n=18 oocytes, 6 frogs), G2019S (n=22 oocytes, 6 frogs) or G2019S+MLi-2 (n=22 oocytes, 6 frogs). Statistical analysis in B was performed using Mann-Whitney test, in E using Kruskal-Wallis test followed by Dunn’s multiple comparisons test. Statistical analysis in F was performed using Unpaired T-test to compare EAAT2+LRRK2 WT to EAAT2+G2019S injected oocytes and with Paired T-test to compare EAAT2+G2019S oocytes before and after LRRK2 inhibition.

We next imaged EAAT2 in oocyte slices to understand the reduction of transport current observed in the presence of LRRK2 G2019S. Fluorescence images and integrated density (IntDen) quantification revealed that the transporter was mainly found at the plasma membrane in oocytes injected with LRRK2 WT, while it was sparsely localized there upon pathogenic LRRK2 G2019S expression (Fig. 3D,E EAAT2+LRRK2 WT vs EAAT2+LRRK2 G2019S, p=0.001). Importantly, short-term incubation of LRRK2 G2019S-injected oocytes with the LRRK2 kinase inhibitor MLi-2 restored both the localization (Fig. 3D-E; EAAT2+LRRK2 G2019S vs EAAT2+LRRK2 G2019S+MLi-2, p=0.0005) and the activity of the transporter (Fig. 3F; EAAT2+LRRK2 G2019S, mean amplitude 17±1 nA vs EAAT2+LRRK2 G2019S+MLi-2 mean amplitude 37±3 nA; p<0.0001; EAAT2+LRRK2 WT: mean amplitude 34±4 vs LRRK2 G2019S+MLi-2; p=0.715). Together with our findings on mouse gliosomes, these observations suggest that the presence of pathogenic human LRRK2 G2019S does not alter the catalytic properties of the glutamate transporter. However, the mutation affects the amount of glutamate transporter functionally trafficked to/from the plasma membrane in a manner mediated by the LRRK2 kinase activity.

### Glt-1 displays an altered localization in Lrrk2 G2019S astrocytes

To gain insights into the mechanism behind the mutant LRRK2-dependent Glt-1 mislocalization and reduced protein levels, we first performed an unbiased protein-protein interaction screen. Endogenous Glt-1 was immunoprecipitated from homogenized WT and Lrrk2 G2019S mouse striata (Supplementary Fig. 2A). As negative control, only magnetic beads were incubated with WT and Lrrk2 G2019S solubilized brain extracts, respectively. Glt-1 interactors were revealed using liquid-chromatography coupled with tandem mass spectrometry (LC-MS/MS) (Supplementary file 1). The two interactome datasets were compared to highlight significant differences in protein abundances (Supplementary file 2). Common and divergent Glt-1 binders between the Lrrk2 WT and G2019S genetic environments were identified upon filtering hits with fold-change >3.5 with respect to the relative control. As shown in the Supplementary Fig. 2B, 40 interactors were present in both datasets, 14 were unique to Glt-1 immunoprecipitated from Lrrk2 WT background and 10 were unique to Lrrk2 G2019S background. Gene Ontology (GO) analysis was employed to identify discrete functional enrichments, and data relative to the Cellular Component (CC) biological domain were analyzed in detail (Supplementary file 3). In both datasets Glt-1 was in contact with membrane domains (GO:0045121; GO:0098857), synaptic and vesicular components (GO:0045202; GO:0043232), cytoskeletal scaffolds (GO:0005856; GO:0005874) and mitochondrial elements (GO:0005743; GO:0031966; GO:0005740) (Supplementary Fig. 2C) as already reported [24]. Components of the plasma membrane region associated with sodium-potassium exchanging ATPase complexes were represented in the Lrrk2 WT genotype only (GO:0098796; GO:0098590; GO:0005890) (Supplementary Fig. 2C). Among the GO terms that characterize the transporter in the Lrrk2 G2019S genotype, we indeed observed several unique categories associated with internalized vesicles and clathrin-mediated endocytosis (GO:0030119; GO:0030128) (Supplementary Fig. 2C). Overall, the pathogenic Lrrk2 G2019S mutation changes the interactome of endogenous striatal Glt-1 toward specific components of the endo-vesicular pathways.

To investigate the localization of Glt-1 in the genotypes, we acutely transfected primary striatal astrocytes with a GFP-Glt-1 construct. TIRFM imaging was performed to selectively visualize Glt-1 localization within thin regions of the plasma membrane [14]. We observed that GFP-Glt-1 localized in or immediately below the plasma membrane in Lrrk2 WT astrocytes, showing a sparse ‘patchy’ pattern (Fig. 4A – WT inset). Although showing a similar membrane distribution, Glt-1 was less detectable on the cell surface of Lrrk2 G2019S astrocytes (Fig. 4A – GS inset). The quantification of Glt-1 mean fluorescence using TIRFM confirmed a lower presence of Glt-1 at the plasma membrane of Lrrk2 G2019S astrocytes as compared to Lrrk2 WT astrocytes (Fig. 4B, Lrrk2 WT vs Lrrk2 G2019S, p<0.0001). However, the application of the Lrrk2 inhibitor MLi-2 in G2019S astrocytes completely rescue the expression of Glt-1 at the plasma membrane, as confirmed by the mean fluorescence quantification (Fig. 4B, Lrrk2 G2019S vs Lrrk2 G2019S+MLi-2, p=0.005; Lrrk2 G2019S+MLi-2 vs Lrrk2 WT p=0.0001). We then studied the overall Glt-1 distribution in primary astrocytes transfected with Flag-Glt-1 using epifluorescence microscopy. Despite the equal ectopic expression of the transporter (Supplementary Fig. 3 A,B: Lrrk2 WT vs Lrrk2 G2019S, p>0.05), we detected a clear accumulation of Glt-1 in round intracellular clusters along with the spotted membrane distribution in Lrrk2 G2019S astrocytes (Fig. 4D). Glt-1 was present within two distinct populations of clusters, namely a larger cluster of ∼1.8 μm diameter and a smaller cluster of ∼0.8 μm diameter, respectively (Supplementary Fig. 3C-F). In contrast, a unique population of dots with an average diameter of 0.6 μm was present in the Lrrk2 WT background, perhaps reflecting clusters in the plasma membrane (Supplementary Fig.3C-D). Using intracellular cluster formation as a readout (diameter >1μm), we detected a significant increase of Glt-1 clusters in Lrrk2 G2019S astrocytes as compared to the Lrrk2 WT genotype (Fig. 4D-E Lrrk2 WT vs Lrrk2 G2019S, p<0.0001). Noteworthy, 90 min MLi-2 application restored the number of Glt-1-positive clusters to control values in Lrrk2 G2019S astrocytes (Fig. 4D-E; Lrrk2 G2019S vs Lrrk2 G2019S+MLi-2, p<0.0001; Lrrk2 G2019S+MLi-2 vs Lrrk2 WT, p>0.99). Conversely, LRRK2 inhibition did not change the number of clusters in Lrrk2 WT astrocytes (Fig. 4D-E Lrrk2 WT vs Lrrk2 WT+MLi-2, p=0.9692). To investigate a possible role for Lrrk2 in PKC-mediated internalization of Glt-1, we used the PKC activator TPA, as well as the PKC inhibitor Go 6976 (Fig. 4C). In agreement with published data [23, 109], the application of TPA enhanced Glt-1 intracellular clusters in Lrrk2 WT astrocytes (Fig. 4D-E; Lrrk2 WT vs Lrrk2 WT+TPA, p<0.0001). Of note, the distribution of cluster diameters in this condition peaked at ∼1.9 μm, which is similar to that observed in Lrrk2 G2019S astrocytes (Supplementary Fig.3 C,F). No significant changes were observed in the number of Glt-1-positive puncta in Lrrk2 G2019S astrocytes upon TPA treatment, suggesting that the majority of Glt-1 was localized to the intracellular compartment already under basal conditions (Fig. 4D-E; Lrrk2 G2019S vs Lrrk2 G2019S+TPA, p=0.9741). Furthermore, to unravel whether LRRK2 acts upstream of PKC, we applied MLi-2 to TPA-treated Lrrk2 WT astrocytes. Interestingly, the application of MLi-2 did not restore the number of Glt-1 clusters to basal values (Fig. 4D-F; Lrrk2 WT vs Lrrk2 WT+TPA+MLi-2, p<0.0001). Moreover, PKC inhibition using Go 6976 did not prevent Glt-1 clustering in Lrrk2 G2019S astrocytes (Fig. 4D-G; Lrrk2 G2019S vs Lrrk2 G2019S+Go 6976, p=0.1645). These data demonstrate that Lrrk2 G2019S reduces Glt-1 localization at the plasma membrane in a PKC-independent manner in primary astrocytes. Moreover, the correction of the pathogenic phenotype observed upon application of MLi-2 indicates that the LRRK2 kinase activity is crucial for the relocation of Glt-1.

**Figure 4.**
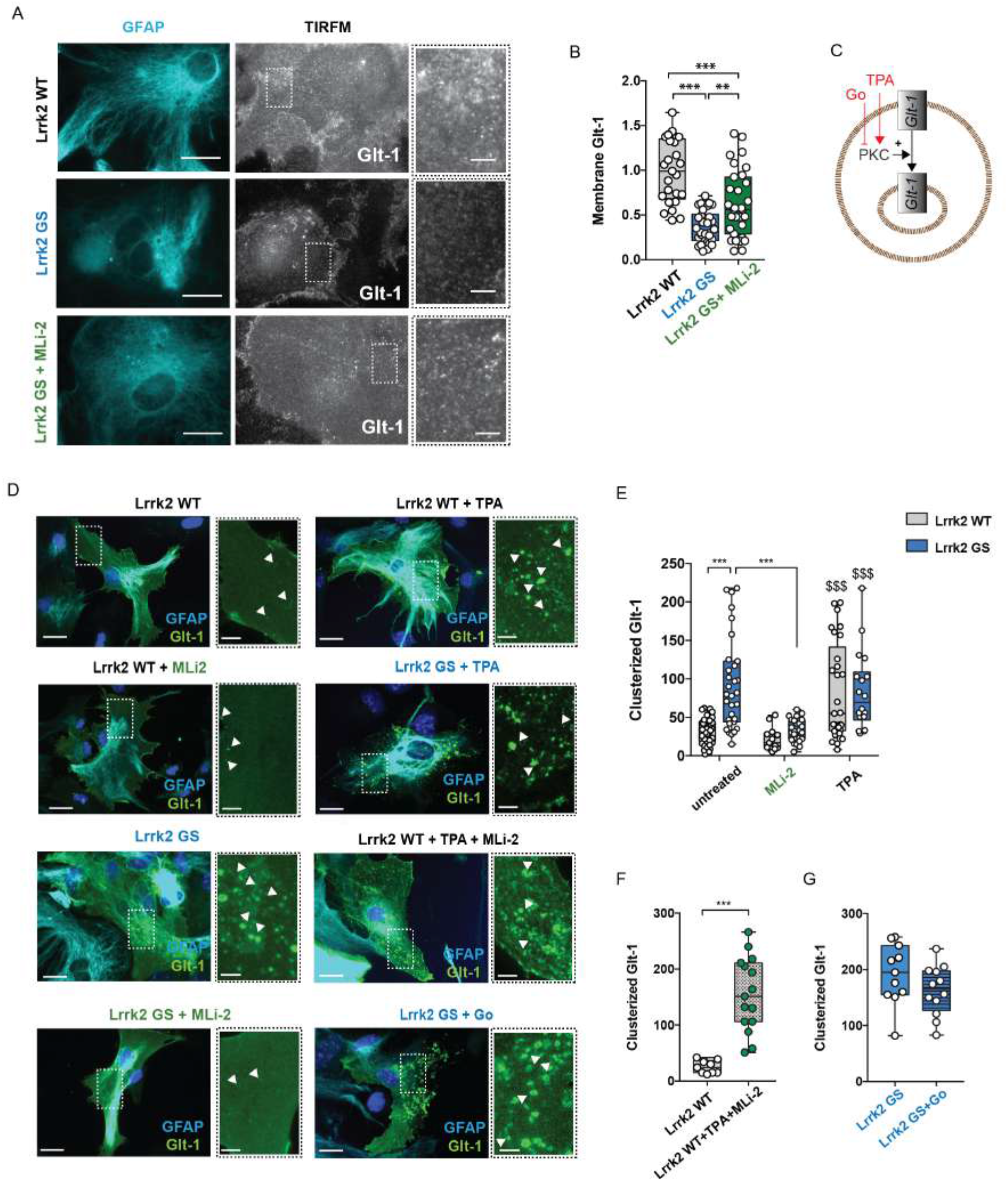
Astroglial Glt-1 is mislocalized in the presence of Lrrk2 G2019S mutation. A) Representative TIRFM images of Glt-1 localization in Lrrk2 WT and Lrrk2 G2019S astrocytes (untreated of treated with MLi-2) transfected with GFP-Glt1 (gray) and stained with GFAP (cyan). Scale bar 20 μm; insets 10 μm; B) Quantification of the GFP-Glt-1 mean fluorescence performed in TIRFM images (n=25 cells from Lrrk2 WT, n=30 cells from Lrrk2 GS and n=31 cells from Lrrk2 GS+MLi-2, experiments performed at least in triple); C) Schematic representation of TPA and Go 6976 effects on PKC activity; D) Representative epifluorescence images of Glt-1 intracellular clusters in Lrrk2 WT and Lrrk2 G2019S astrocytes transfected with Flag-Glt-1 (green) and stained with GFAP (cyan) under basal condition and after pharmacological treatment; Scale bar 20 μm, insets 5 μm; E) Quantification of the number of Glt-1-positive clusters per cell in Lrrk2 WT and G2019S astrocytes under basal condition and after pharmacological treatment using Lrrk2 inhibitor MLi-2 (90 min) and the PKC activator TPA (20 min). Number cells analyzed: Lrrk2 WT (n= 40 cells), Lrrk2 WT+MLi-2 (n= 13 cells), Lrrk2 G2019S (n=31 cells), Lrrk2 G2019S+MLi-2 (n=26 cells), Lrrk2 WT+TPA (n=29 cells), Lrrk2 G2019S+TPA (n=18 cells); F) Quantification of the number of Glt-1-positive clusters per cell under basal condition and after pharmacological co-treatment with TPA and Mli-2 in Lrrk2 WT astrocytes. Number cells analyzed: Lrrk2 WT (n= 9 cells) and WT+TPA+MLI-2 (n=15 cells); G) Quantification of the number of Glt-1-positive clusters per cell under basal condition and after pharmacological treatment with the PKC inhibitor Go 6976 in Lrrk2 G2019S astrocytes. Number cells analyzed: Lrrk2 GS (n= 11 cells) and Lrrk2 GS+Go (n=12 cells). Experiments performed at least in triple. Three cells analyzed for each independent cell cultures. Statistical analysis in B was performed using One-way ANOVA test followed by Tukey’s multiple comparisons test. Statistical analysis in E was performed using Two-way ANOVA test followed by Tukey’s multiple comparisons test ($ vs Lrrk2 WT and * versus Lrrk2 G2019S). Statistical analysis in F and G was performed using Unpaired T-test.

### Glt-1 is retained at the Rab4-positive compartment in Lrrk2 G2019S striatal astrocytes

To determine the identity of the Glt-1 clusters, we co-transfected Lrrk2 WT and Lrrk2 G2019S astrocytes with Flag-Glt-1 together with GFP-Lamp1 (lysosomal marker), GFP-Rab11 (marker of slow-recycling endocytosis) or GFP-Rab4 (marker of fast-recycling endocytosis) (Fig. 5A). Quantification showed that there was no significant difference in the amount of Lamp1, Rab11 or Rab4 fluorescence between the two genotypes (Fig. 5B,D,F; all comparisons p>0.05). We then quantified the co-localization of Glt-1 with the three markers. There were no significant changes in the amount of the transporter in the Lamp1- or Rab11-positive compartments in Lrrk2 G2019S astrocytes as compared to Lrrk2 WT astrocytes (Fig. 5C,E; all comparisons p>0.05). However, there was a significant increase in the amount of Glt-1 co-localizing with the Rab4-positive fast-recycling vesicles in Lrrk2 G2019S astrocytes as compared to control (Fig. 5G, Lrrk2 WT vs Lrrk2 G2019S, p=0.001). Interestingly, short-term application of MLi-2 reverted the Lrrk2 G2019S-mediated pathogenic phenotype as shown by the loss of co-localization between Glt-1 and Rab4-positive vesicles (Fig. 5G, Lrrk2 G2019S vs Lrrk2 G2019S+MLi-2, p=0.001; Lrrk2 G2019S +MLi-2 vs Lrrk2 WT, p=0.98). Z-stack orthogonal projections confirmed that Glt-1 co-localized with Rab4-positive vesicles in Lrrk2 G2019S primary astrocytes (Supplementary Fig. 4A).

**Figure 5.**
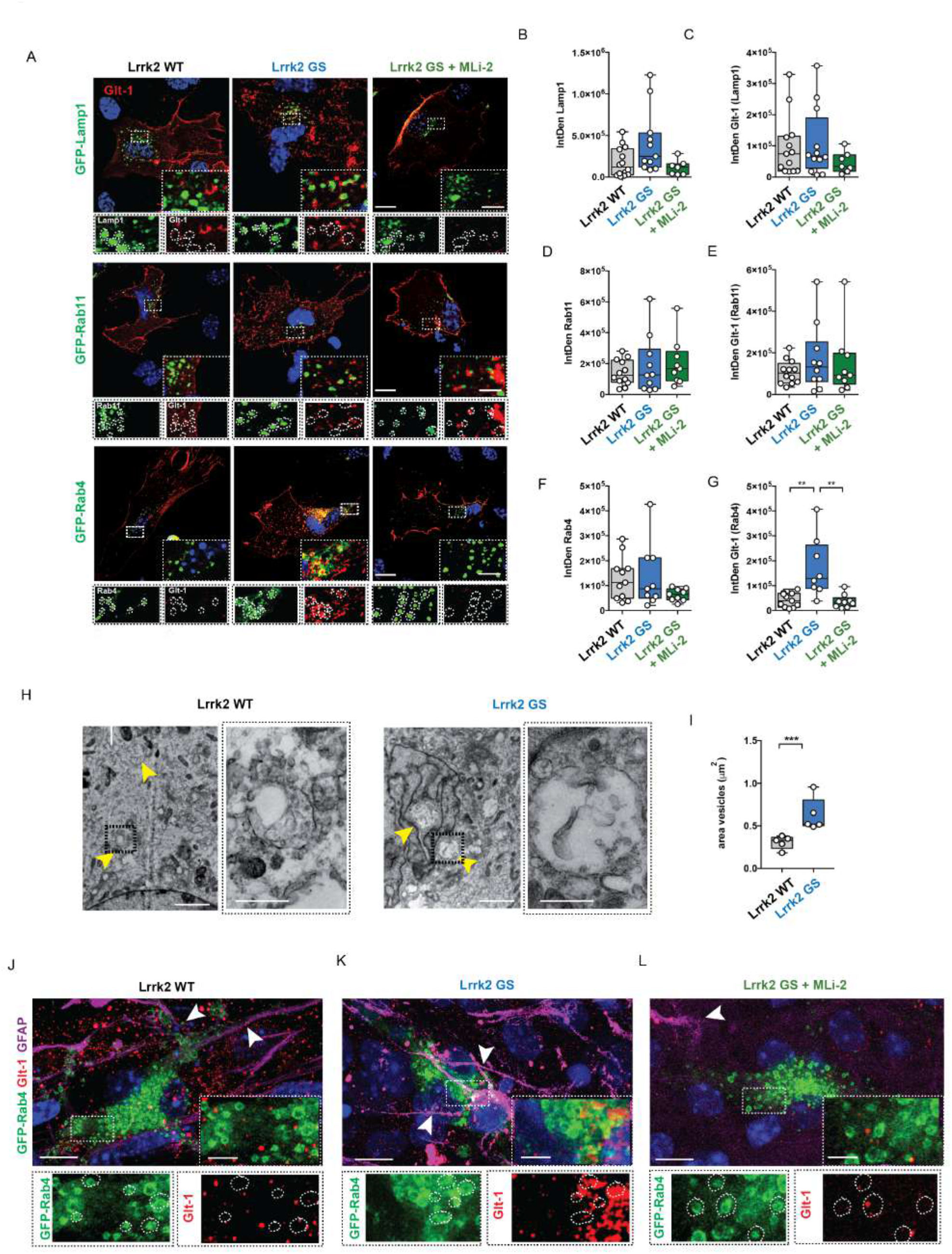
Lrrk2 G2019S enhances Glt-1 accumulation in Rab4-positive organelles. A) Representative z-stack confocal images of primary astrocytes from Lrrk2 WT and Lrrk2 G2019S mice transfected with Flag-Glt-1 and GFP-Lamp1, GFP-Rab11 or GFP-Rab4 under basal conditions or upon treatment with MLi-2 inhibitor. Insets show Lamp1-, Rab11- or Rab4-positive area and the localization of Glt-1 in the indicated ROIs. Scale bars: 20 μm; insets 5 μm; B-C) Quantitative analysis of Lamp1 IntDen (n=14 cells for Lrrk2 WT, n=11 cells for Lrrk2 GS and n=7 cells for Lrrk2 GS+MLi2, experiments performed at least in triple) and Glt-1 IntDen in Lamp1 compartment (n=13 cells for Lrrk2 WT, n=12 cells for Lrrk2 GS and n=7 cells for Lrrk2 GS+MLi2, experiments performed at least in triple); D-E) Quantitative analysis of Rab11 IntDen and Glt-1 IntDen in Rab11 compartment (n=12 cells for Lrrk2 WT, n=10 cells for Lrrk2 GS and n=9 cells for Lrrk2 GS+MLi2, experiments performed at least in triple); F-G) Quantitative analysis of Rab4 IntDen (n=11 cells for Lrrk2 WT, n=9 cells for Lrrk2 GS and n=9 cells for Lrrk2 GS+MLi2, experiments performed at least in triple) and Glt-1 IntDen in Rab4 compartment (n=11 cells for Lrrk2 WT, n=8 cells for Lrrk2 GS and n=9 cells for Lrrk2 GS+MLi2, experiments performed at least in triple). H) Representative TEM images of Lrrk2 WT and G2019S endosomal-like structures (yellow arrowheads) in primary striatal astrocytes transfected with GFP-Rab4 and Flag-Glt-1 (scale bars: 5 μm; insets: scale bars: 500 nm); I) Quantification of the area of endosomal-like structures (n=5 cells analyzed for each group, ten independent fields were analyzed for quantification). J-L) Representative z-stack confocal images of coronal organotypic Lrrk2 WT, Lrrk2 G2019S and Lrrk2 G2019S treated with MLi-2 slices transfected with the GFP-Rab4 plasmid and stained for the endogenous proteins Glt-1 (red) and GFAP (magenta). Insets show Rab4-positive area and the localization of Glt-1 in the indicated ROIs. Scale bars: 20 μm; insets 2 μm; Statistical analysis in B-G was performed using One-way ANOVA test followed by Tukey’s multiple comparison test. Statistical analysis in I was performed using Unpaired t-tests.

Since pathogenic LRRK2 did not influence the expression of Rab4 protein *per se*, we investigated the impact of the kinase on Rab4-positive vesicle morphology using both confocal fluorescence microscopy and TEM (Supplementary Fig. 4B-E and Fig.5H). Fluorescence analysis demonstrated that the Lrrk2 G2019S mutation did not alter the total amount of Rab4-positive vesicles (Supplementary Fig. 4 B,C, Lrrk2 WT vs Lrrk2 G2019S, p>0.05). Rather, the area of Rab4-positive fast recycling endosomes appeared increased in the presence of the pathogenic Lrrk2 mutation (Supplementary Fig.4 B, D, Lrrk2 WT vs Lrrk2 G2019S, p=0.003). By overlapping fluorescence images with TEM ultrathin sections, we were able to identify and define the Rab4-positive recycling vesicles (Supplementary Fig.4E-asterisks). Rab4-positive endosomes presented an irregular shaped vacuole (0.2-0.5 μm diameter) characterized by an electron-lucent lumen [19, 41]. TEM analysis confirmed that the Lrrk2 G2019S mutation markedly increased the area of Rab4-positive vesicles (Fig. 5H,I; Lrrk2 WT vs Lrrk2 G2019S, p<0.0001) as compared to controls.

To validate these observations in intact tissue, we employed organotypic coronal cultures derived from Lrrk2 WT and Lrrk2 G2019S mice. Slices were transfected with the GFP-Rab4 plasmid to visualize the fast-recycling compartment. As shown in Fig. 5J-L, the endogenous expression of Glt-1 was reduced in Lrrk2 G2019S organotypic slices as compared to control samples. Moreover, astrocytic endogenous Glt-1 abundantly accumulated in Rab4-positive vesicles in the presence of the G2019S mutation (Fig. 5K) compared to the WT counterpart (Fig. 5J). A representative 3D image confirming the Glt-1 colocalization with Rab4-positive vesicles in the pathogenic compartment is shown in Supplementary Files 4. Intriguingly, short-term application of the Lrrk2 kinase inhibitor MLi-2 restored the phenotype (Fig. 5L). Overall, here we show that in Lrrk2 G2019S astrocytes the transporter is predominantly confined to enlarged Rab4-positive recycling vesicles not only in primary astrocytes but also in organotypic slices. In both cases, the rescuing effect of MLi-2 suggests that dysregulation of the Lrrk2 kinase activity is sufficient to impair Glt-1 trafficking.

### G2019S Lrrk2 alters the endocytic Rab4-positive compartment *via* Rab8A/Rab10

Multiple reports indicate that LRRK2 regulates recycling and secretory trafficking pathways *via* the phosphorylation and inactivation of its substrates Rab8A and Rab10 [33, 72, 74]. We first evaluated the levels and activity of endogenous Lrrk2 in WT and G2019S Lrrk2 astrocytes by monitoring two different phosphosites: pS935 (CK1α/PKA/IKKs phosphorylation site), used to detect on-target effects of LRRK2 kinase inhibitor treatment, and pS1292 (auto-phosphorylation site), used to directly measure LRRK2 kinase activity (Fig. 6A). In agreement with previous reports, G2019S Lrrk2 decreased the levels of pS935-Lrrk2 [34, 64, 101] and increased the levels of pS1292-Lrrk2 [34, 100, 101] (Fig. 6A). Total Rab8A and Rab10 levels were not different between genotypes, whilst pT72-Rab8A and pT73-Rab10 levels were increased in the G2019S Lrrk2 astrocytes (Fig. 6A). As expected, acute application of the Lrrk2 kinase inhibitor MLi-2 to G2019S Lrrk2 astrocytes reverted the increased phosphorylation levels of Rab8A, Rab10 and pS1292-Lrrk2, and abolished the levels of pS935-Lrrk2 (Fig. 6A).

**Figure 6.**
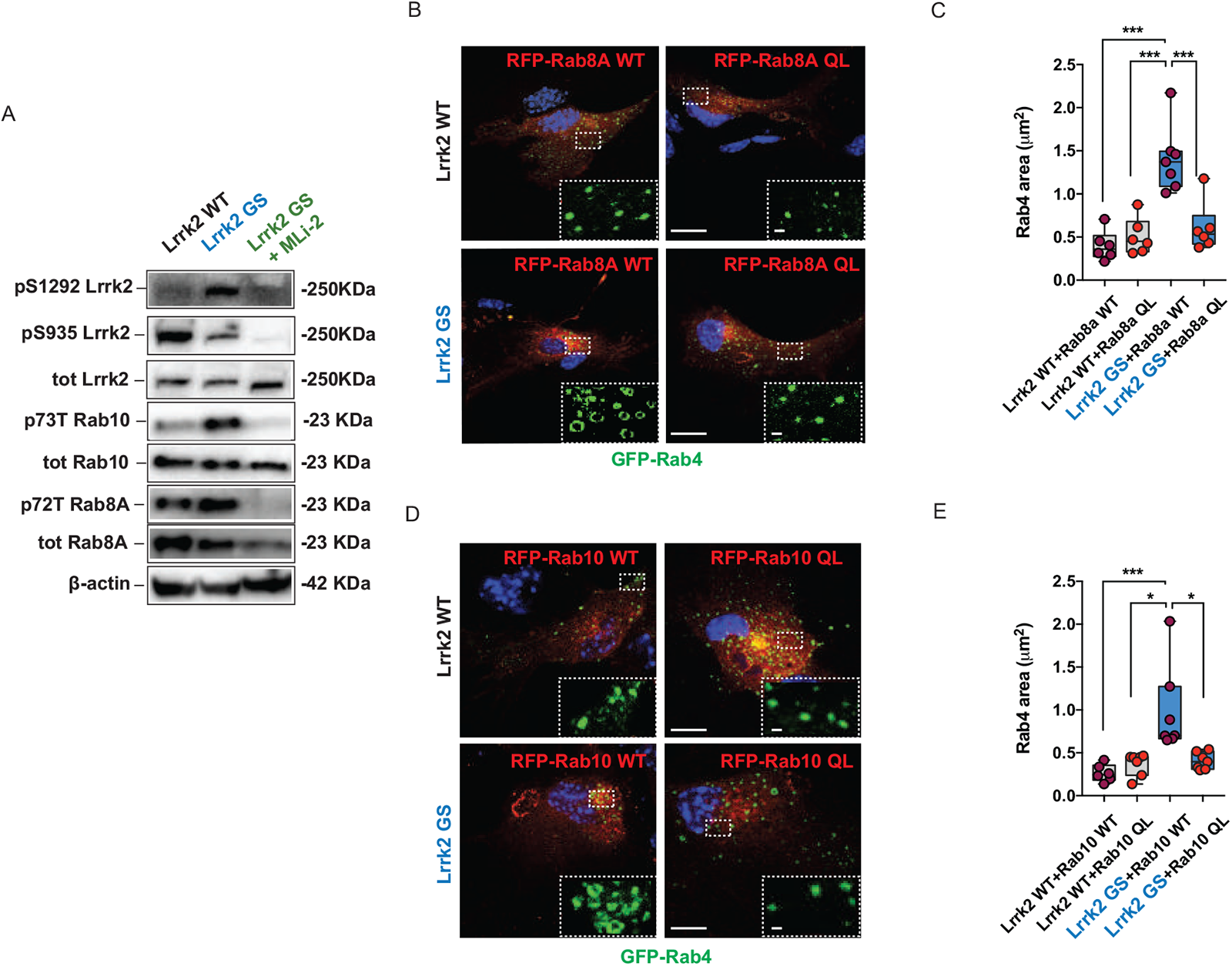
Lrrk2 G2019S mutation enhances Rab4-positive organelles dimension via Rab8A and Rab10 phosphorylation. A) Western blot analysis of primary striatal Lrrk2 WT, Lrrk2 G2019S and Lrrk2 G2019S treated with the Lrrk2 inhibitor MLi-2 astrocytes using anti-pS1292 Lrrk2, anti-pS935 Lrrk2, anti-total Lrrk2, anti-p73 Rab10, anti-total Rab10, anti-p72 Rab8A and anti-total Rab8A antibodies; B,D) Representative z-stack confocal images of primary astrocytes from Lrrk2 WT and Lrrk2 G2019S mice transfected with GFP-Rab4 and Rab8A WT/QL or Rab10 WT/QL. Insets show the area of the Rab4-positive structures. Scale bars: 20 μm; insets 1 μm; C,E) Quantification of Rab4-positive vesicle area in primary astrocytes from Lrrk2 WT and Lrrk2 G2019S mice transfected with GFP-Rab4 and Rab8A WT/QL or Rab10 WT/QL; at least n=6 cells have been analyzed for each experimental group and the experiments have been performed in triple. Statistical analysis in C was performed using One-way ANOVA test followed by Tukey’s multiple comparisons test; Statistical analysis in E was performed using Kruskal-Wallis test followed by Dunn’s multiple comparisons test.

To test whether the Lrrk2 -mediated alterations of the Rab4-positive recycling compartment are modulated by Rab8A or Rab10, we transfected primary astrocytes with Flag-Glt-1 and GFP-Rab4 along with either RFP-tagged wild type Rab8A or Rab10, or with variants mimicking the GTP-bound active state (Rab8A-Q67L or Rab10-Q68L) (Fig. 6B,D). Overexpression of wildtype Rab8A or wildtype Rab10 did not alter the G2019S Lrrk2-mediated increase in the size of the Rab4-positive recycling endosomes (Fig. 6 C-E; Lrrk2 WT+Rab8A WT vs Lrrk2 G2019S+Rab8A WT, p<0.0001; Lrrk2 WT+Rab10 WT vs Lrrk2 G2019S+Rab10 WT, p=0.0003). In contrast, overexpression of the active variants of RFP-Rab8A or RFP-Rab10 completely reverted the G2019S Lrrk2-mediated morphological alterations of the Rab4 recycling compartment (Fig. 6 C-E; Lrrk2 G2019S+Rab8A-Q67L vs Lrrk2 WT+Rab8A-Q67L, p=0.91; Lrrk2 G2019S+Rab8A-Q67L vs Lrrk2 G2019S+Rab8A WT, p=0.0003; Lrrk2 G2019S+Rab10-Q68L vs Lrrk2 WT+Rab10-Q68L, p>0.99; Lrrk2 G2019S+Rab10-Q68L vs Lrrk2 G2019S+Rab10 WT, p=0.04). Together, analyzing the distribution of Rab4-positive vesicles dimension (Supplementary Fig.5 A-B), we confirmed that the overexpression of Rab8A-Q67L or Rab10-Q68L variants in Lrrk2 G2019S astrocytes shifted vesicle dimension range from ∼1-0.8 μm^2^ to ∼0.5-0.2 μm^2^, sizes similar to wildtype organelles (Supplementary Fig.5 A-B). Altogether, these data suggest that the aberrant kinase activity of pathogenic Lrrk2 causes enlargement of the Rab4-positive compartment through Rab8A and Rab10 inactivation.

### Lrrk2 G2019S perturbs Glt-1 recycling and favors its degradation

We next examined whether Glt-1 accumulation in the Rab4-positive compartment in the Lrrk2 G2019S astrocytes was due to a delay in recycling. First, we applied the recycling inhibitor Monensin on primary striatal Lrrk2 WT and G2019S astrocytes and investigated the co-localization of Glt-1 with Rab4 (Fig. 7A). In untreated cells, Glt-1 accumulated in Rab4-positive recycling vesicles in the presence of the Lrrk2 pathogenic mutation as compared to controls (Fig. 7B; untreated Lrrk2 WT vs untreated Lrrk2 G2019S, p=0.003). Monensin application induced a significant accumulation of Glt-1 in Rab4-positive vesicles in Lrrk2 WT cells (Fig. 7B; untreated Lrrk2 WT vs Lrrk2 WT+Monensin, p=0.001), thereby phenocopying the distribution of the transporter observed in untreated Lrrk2 G2019S astrocytes (Fig. 7B; Lrrk2 WT+Monensin vs untreated Lrrk2 G2019S, p>0.99). Conversely, Monensin treatment did not induce further Glt-1 accumulation in Rab4-positive vesicles in the Lrrk2 G2019S astrocytes (Fig. 7B; untreated Lrrk2 G2019S vs Lrrk2 G2019S+Monensin, p>0.99), suggesting that the transporter was almost completely incorporated in the fast-recycling compartment under basal conditions in those cells.

**Figure 7.**
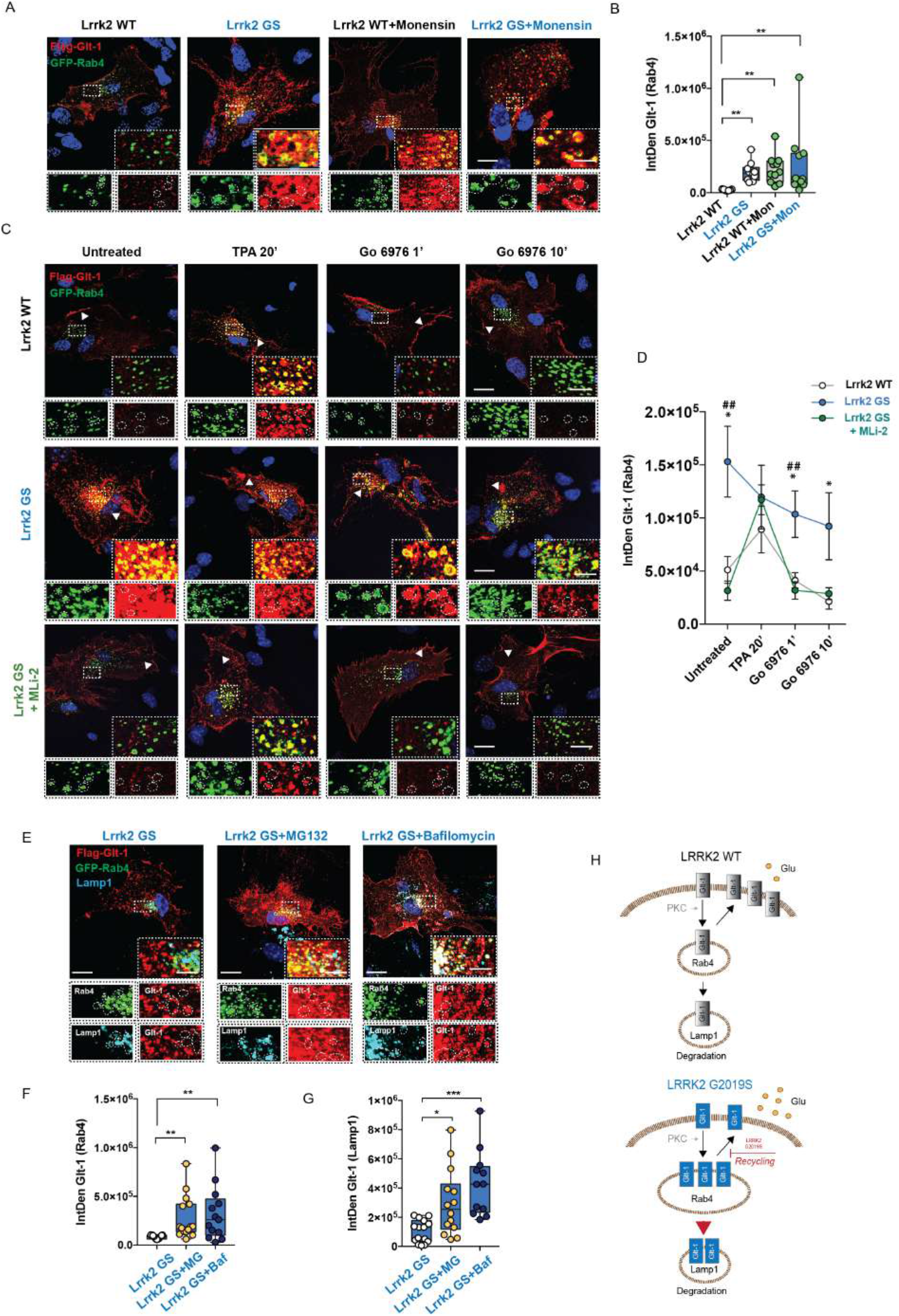
G2019S pathogenic Lrrk2 mutation impacts on Glt-1 recycling and turnover. A) Representative confocal images of primary striatal Lrrk2 WT and G2019S astrocytes under basal conditions and treated with the recycling blocker Monensin (35 µM, 40 min). Insets show the Rab4-positive area and the localization of Glt-1 in the indicated ROIs. Scale bars: 20 μm; insets 5 μm; B) Quantitative analysis of Glt-1 IntDen in Rab4-positive compartment; n= 7 cells for Lrrk2 WT, n=10 cells for Lrrk2 WT+Monensin, n=11 cells for Lrrk2 G2019S and n=11 cells for Lrrk2 G2019S +Monensin, experiments performed at least in triple; C) Representative z-stack confocal images of primary Lrrk2 WT and G2019S (with or without MLi-2) striatal astrocytes transfected with Flag-Glt-1 (red) and GFP-Rab4 (green). The insets show Rab4-positive area and the localization of Glt-1 in these ROIs; scale bar 20 μm; insets 5 μm; D) Quantification of Glt-1 IntDen in Rab4 compartment in the four selected experimental time points. At least n=6 cells have been analyzed for each experimental group and the experiments have been performed in triple; E) Representative z-stack confocal images of primary striatal of Lrrk2 G2019S astrocytes transfected with GFP-Rab4 and Flag-Glt-1 (red) and labeled for endogenous Lamp1 (far red, pseudocolored in blue). Insets show Rab4- and Lamp1-positve area and the localization of Glt-1 in these ROIs. Scale bar 20 μm; insets 5 μm; F) Quantification of Glt-1 IntDen in the Rab4-positive vesicles in Lrrk2 G2019S astrocytes under basal conditions (n=13 cells) or upon MG132 (n=15 cells) or Bafilomycin application (n=13 cells); G) Quantification of Glt-1 IntDen in the endogenous Lamp1-positive structures in Lrrk2 G2019S astrocytes under basal conditions (n=15 cells) or upon MG132 (n=14 cells) or Bafilomycin (n=12 cells) application; experiments performed at least in triple. H) Schematic representation of Glt-1 trafficking in Lrrk2 WT and Lrrk2 G2019S astrocytes. Statistical analysis in B,F was performed using Kruskal-Wallis test followed by Dunn’s multiple comparisons test. Statistical analysis in D and G was performed using One-way ANOVA test followed by Tukey’s multiple comparisons test (* Lrrk2 WT vs Lrrk2 G2019S, ## Lrrk2 G2019S vs Lrrk2 G2019S+MLi-2).

We next studied the recycling kinetics by measuring Glt-1 localization in Rab4-positive vesicles after TPA-induced internalization. To isolate the unique contribution of the recycling pathway, the subsequent basal internalization was blocked using Go 6976. We again confirmed that under basal conditions Glt-1 accumulated as clusters in Rab4-positive vesicles in Lrrk2 G2019S but not in Lrrk2 WT astrocytes (Fig. 7C-D; Lrrk2 WT vs Lrrk2 G2019S, p=0.01), and that the application of MLi-2 reverted this phenotype (Fig. 7C-D; Lrrk2 G2019S vs Lrrk2 G2019S+MLi-2, p=0.002; Lrrk2 G2019S+MLi-2 vs Lrrk2 WT, p= 0.79). Upon TPA stimulation, there was an overall increase of Glt-1 in Rab4-positive vesicles in Lrrk2 WT and Lrrk2 G2019S+MLi-2 astrocytes, reaching values of Glt-1/Rab4 co-localization similar to those observed in Lrrk2 G2019S astrocytes (Fig. 7C-D; Lrrk2 WT+TPA vs Lrrk2 G2019S+TPA, p=0.63; Lrrk2 G2019S+TPA vs Lrrk2 G2019S+MLi-2+TPA, p=0.99; Lrrk2 WT+TPA vs Lrrk2 G2019S+MLi-2+TPA, p=0.70). Conversely, the amount of Glt-1 co-localizing with Rab4-positive vesicles in Lrrk2 G2019S astrocytes did not increase upon TPA stimulation (Fig. 7C-D). Both Lrrk2 WT and Lrrk2 G2019S+MLi-2 astrocytes displayed a significant decrease of Glt-1 in Rab4-positive vesicles already 1 min after Go 6976 application (Fig. 7C-D; Lrrk2 WT+ Go T1’ vs Lrrk2 G2019S+MLi-2+Go T1’, p=0.88; Lrrk2 WT+ Go T1’ vs Lrrk2 WT, p=0.95; Lrrk2 G2019S+MLi-2+Go T1’ vs Lrrk2 G2019S+MLi-2, p>0.99). In contrast, Glt-1 was almost entirely retained in Rab4-positive vesicles in Lrrk2 G2019S astrocytes at this time point (Fig. 7C-D; Lrrk2 WT+ Go T1’ vs Lrrk2 G2019S+Go T1’, p=0.01; Lrrk2 G2019S+MLi-2+Go T1’ vs Lrrk2 G2019S+Go T1’, p=0.007; Lrrk2 G2019S+Go T1’ vs Lrrk2 G2019S, p=0.64). A most prominent effect was observed 10 min after Go 6976 application (Fig. 7C-D; Lrrk2 WT+ Go T10’ vs Lrrk2 G2019S+Go T10’, p=0.03; Lrrk2 G2019S+MLi-2+Go T10’ vs Lrrk2 G2019S+Go T10’, p=0.06; Lrrk2 WT+ Go T10’ vs Lrrk2 G2019S+MLi-2+Go T10’, p=0.95; Lrrk2 WT+ Go T10’ vs Lrrk2 WT, p=0.39; Lrrk2 G2019S+MLi-2+Go T10’ vs Lrrk2 G2019S+MLi-2, p=0.99). Although not reaching statistical significance, the overall amount of the transporter at the Rab4-positive organelles progressively decreased in the pathogenic Lrrk2 astrocytes upon Go treatment (Fig. 7C-D; Lrrk2 G2019S+ Go T10’ vs Lrrk2 G2019S, p=0.49). Of note, this phenomenon was not associated with a relocation of Glt-1 to the plasma membrane in the G2019S background (Fig. 7C, arrowheads).

Therefore, we examined whether Glt-1 incorporated into Rab4-positive vesicles was targeted for degradation in the Lrrk2 G2019S astrocytes. As Glt-1 can be degraded through both the proteasome [80, 99] and the lysosome [88, 107], we monitored the presence of Glt-1 in the fast recycling and lysosomal compartments in the presence of either MG132 or Bafilomycin to block proteasomal or lysosomal function, respectively [67, 80]. Lrrk2 G2019S astrocytes were co-transfected with Flag-Glt-1 and GFP-Rab4 and stained for the endogenous lysosomal protein Lamp1 to measure Glt-1 localization in the Rab4- and Lamp1-positive organelles (Fig. 7E). Upon MG132 application, the fraction of Glt-1 co-localizing with the Rab4-positive compartment was significantly enhanced in Lrrk2 G2019S astrocytes (Fig. 7F; Lrrk2 G2019S vs Lrrk2 G2019S+MG132; p=0.0012). In the same condition, MG132 also promoted increases of Glt-1 localization in the Lamp1-positive vesicles in Lrrk2 G2019S astrocytes (Fig. 7G; Lrrk2 G2019S vs Lrrk2 G2019S+MG132; p=0.0196). Furthermore, we demonstrated that lysosomal inhibition by Bafilomycin caused a significant increase of Glt-1 in the Rab4-positive compartment (Fig. 7F; Lrrk2 GS and Lrrk2 G2019S+Bafilomycin; p=0.0038) and dramatically accentuated Glt-1 accumulation in Lamp1-positive vesicles in Lrrk2 G2019S astrocytes (Fig. 7G; Lrrk2 G2019S vs Lrrk2 G2019S+Bafilomycin; p=0.0001). Similar accumulation of Glt-1 upon pharmacological manipulations was measured in the wild-type background confirming the involvement of both the degradative pathways in Glt-1 degradation (Supplementary Fig.6 A-B). These results indicate that pathogenic Lrrk2 profoundly perturbs Glt-1 recycling to the cell membrane, possibly associated with the re-routing of the transporter for lysosomal degradation (Fig. 7H).

## Discussion

In the present study, we report severe deficits of EAAT2 in *post-mortem* caudate and putamen from Parkinson’s disease (PD) patients carrying the LRRK2 G2019S mutation. Previous clinical data have shown alterations in the glutamate content in the brain as well as in the plasma of idiophatic PD (iPD) patients, suggesting that glutamate dyshomeostasis might be a common feature in PD [20, 29, 37, 63, 104]. However, our data indicate that the EAAT2 phenotype is clearly exacerbated in the presence of the LRRK2 pathogenic mutation as compared to iPD, suggesting a specific involvement of LRRK2 in EAAT2 biology. Striatal EAAT2 downregulation correlated with increased expression of the reactive marker GFAP in caudate and putamen glial cells of LRRK2-linked PD patients. Although impaired glutamate buffering is often found accompanied by astroglial reactivity and neuroinflammation in several disease models [31, 59, 110], conflicting results have been published in the context of PD patients. Enhanced GFAP reactivity has been clearly documented in *Substantia Nigra pars compacta (SNpc)* and olfactory bulbs [21, 54, 94]. However, one report suggested astrocytic atrophy (instead of reactivity) in PD brains, since low levels of astrocyte markers were observed in the *SNpc* and the striatum [94], and studies in human iPS-derived astrocytes from LRRK2 G2019S-PD patients suggested a similar mechanism [70]. Our study suggests an absence of astrocytic degeneration or atrophy since the levels of the astrocyte marker GS were not perturbed in the context of disease. The presence of astrocyte reactivity has been reported in different transgenic mice overexpressing the LRRK2 G2019S kinase [34, 105]. Accordingly, LRRK2 G2019S knock-in mice display an evident phenotype characterized by a significant downregulation of striatal Glt-1 protein levels and extensive astrogliosis, which recapitulates the human disease in *post-mortem* samples. Thus, the knock-in G2019S LRRK2 mouse is an outstanding model system to unravel the mechanism that links pathogenic LRRK2 to transporter deficiencies. Glt-1 deficits in mice appear without any nigral degeneration, indicating that LRRK2-mediated glutamate transporter dysfunction in the striatum might anticipate dopaminergic cell loss. In agreement with the downregulation of Glt-1 protein, an enhanced glutamatergic cortico-striatal neurotransmission has been described in this animal model[8, 46, 52, 53, 95, 98] These findings suggest that striatal glutamatergic imbalance and correlated glutamate-induced excitotoxicity intervene in PD pathophysiology.

LRRK2 is expressed in glial cells and plays a relevant role in astrocyte physiology both in humans and mice [5, 57, 87]. In this context, we specifically dissected the contribution of the pathogenic LRRK2 mutation to Glt-1 functionality in astrocytes at endogenous levels in cultured cells, in pure subcellular astrocytic *ex vivo* preparations and in organotypic coronal slices. We confirmed a reduced level of the transporter, in agreement with our observations in human and murine striatal lysates. These observations fit with the functional analysis carried out on isolated mouse striatal gliosomes, which reveal a decreased ability of the transporter to reuptake glutamate. By comparing the kinetic values, we find that LRRK2 G2019S impinges on Glt-1 transport velocity without affecting substrate affinity. Similarly, the expression of human LRRK2 G2019S induces a decrease of the EAAT2 transport current without modifying the biophysical properties of the transporter in *Xenopus* oocytes. Therefore, the EAAT2/Glt-1 functional impairment can be ascribed to a reduced localization of the transporter at the plasma membrane mediated by pathogenic LRRK2. Importantly, the effects of G2019S LRRK2 on EAAT2 are restored upon acute inhibition of the G2019S kinase activity by MLi-2, indicating that pathogenic LRRK2 impairs proper EAAT2 homeostasis.

It has been consistently reported that Glt-1 continuously exchanges between a diffuse and spotted localization at the surface, shaping synaptic transmission [2, 61]. Moreover, Glt-1 undergoes basal PKC-mediated endocytosis and intracellular cluster formation in astrocytic processes to regulate glutamate uptake activity [109]. In our system, only the amount of the transporter, but not the spotted distribution at the plasma membrane, was affected by pathogenic LRRK2. Conversely, Lrrk2 G2019S astrocytes exhibited the presence of large intracellular Glt-1 clusters resembling, in terms of dimension, those found upon pharmacological stimulation of PKC-mediated endocytosis of Glt-1. However, our data exclude the involvement of Lrrk2 in the PKC-mediated internalization pathway. Indeed, the inhibition of Lrrk2 kinase activity was not able to revert the PKC-induced clusterization in control astrocytes, and PKC inhibition failed to reduce Glt-1 clusters in G2019S Lrrk2 astrocytes. PKC activation was also not able to induce a further increase in Glt-1 clusters in Lrrk2 G2019S astrocytes, suggesting that the majority of the transporter is internalized in the pathogenic Lrrk2 background. Endogenous Glt-1 interacts with partially different protein environments in the two backgrounds, as revealed by mass spectrometry data. On the one hand, striatal Glt-1 immunoprecipitated from Lrrk2 WT mice mainly interacts with plasma membrane proteins. In agreement with previous findings, we identified the Na^+^-K^+^ pump as a Glt-1 interactor in a Lrrk2 WT background, confirming that Glt-1 and Na^+^-K^+^-ATPases are part of the same macromolecular complex and operate as a functional unit to regulate physiological glutamatergic neurotransmission [76]. On the other hand, Glt-1 almost exclusively interacts with intracellular proteins in presence of the G2019S Lrrk2 mutation. Indeed, we identified proteins involved in the endo-vesicular pathway, suggesting a different subcellular compartmentalization of the transporter in the presence of the Lrrk2 G2019S mutation. Along these lines, we demonstrate that LRRK2 operates in Glt-1 recycling to the plasma membrane. In the Lrrk2 G2019S astrocytes, Glt-1 is preferentially engulfed in fast-recycling endosomes, which are recognized by overexpression of the marker Rab4. Pharmacological LRRK2 inhibition reduces the amount of Glt-1 co-localizing with the Rab4-positive marker and promotes Glt-1 redistribution to the plasma membrane, which allows for the functional recovery of the transporter as assessed by electrophysiological recordings in oocytes. These findings have been corroborated in a more complex and robust physiological system represented by the *ex vivo* organotypic coronal slices. Here, endogenous Glt-1 accumulates in enlarged Rab4-positive vesicles and the acute application of the Lrrk2 inhibitor MLi-2 reverts this phenotype.

By combining confocal and TEM ultrastructural analysis, we show that the Lrrk2 G2019S mutation profoundly affects the architecture of the fast-recycling Rab4-positive organelles in astrocytes, promoting an enlargement of the area of these vesicles. Coherently with previous studies [34, 74, 101], we report that G2019S Lrrk2 aberrantly phosphorylates Rab8A/Rab10 proteins at the endogenous level in cultured astrocytes. The Rab hyperphosphorylation has been reported to promote defects in the recycling of the EGFR and its accumulation in Rab4-positive recycling structures [72, 74]. In line with these reports, we observed that the overexpression of the active Rab8A or Rab10 mutants in Lrrk2 G2019S astrocytes restored the dimension of the Rab4-positive vesicles, pointing to the Lrrk2-mediated phosphorylation and inactivation of Rab8A/Rab10 as the pathogenic mechanism causing the Glt-1 trafficking defects in G2019S Lrrk2 astrocytes. The role of Lrrk2 in Glt-1 recycling was further dissected by pharmacological approaches. The accumulation of Glt-1 in the Rab4-positive compartment observed in Lrrk2 G2019S astrocytes was phenocopied by blocking the recycling of Glt-1 in control astrocytes. In Lrrk2 G2019S astrocytes, the same treatment did not induce further increases in Glt-1 co-localization with Rab4 vesicles, confirming that almost the totality of the transporter is localized to this subcellular compartment under basal conditions. Moreover, our observations on the kinetics of the process confirm that when internalized, Glt-1 is re-routed to early endosomes, as previously reported [27]. While Glt-1 repopulates the plasma membrane in Lrrk2 wild-type astrocytes, Glt-1 is basally captured and persists in the Rab4 compartment in the pathogenic Lrrk2 context. In addition, a tendential decrease in the amount of Glt-1 colocalizing with the Rab4-positive marker not associated with plasma membrane re-localization indicates that the persistent delay in Glt-1 recycling might promote the degradation of the protein. Accordingly, the blockade of the degradative systems promoted an increase in the amount of Glt-1 colocalizing with a Lamp1-positive compartment especially upon lysosomal inhibition.

In conclusion, our work reveals that the LRRK2 G2019S mutation profoundly affects Glt-1 recycling to the plasma membrane by impinging on the early endosomal fast recycling compartment in striatal astrocytes. Although additional mechanistic investigations are needed, we propose that the cellular degradative system may eventually promote Glt-1 turnover by sensing the overload of the transporter in the Rab4-positive compartment (Fig. 7H). In this manner, a chronic, subtle LRRK2-mediated impairment of the recycling machinery in brain cells might cause a progressive depletion of Glt-1 with a consequent reduction of extracellular glutamate clearance. In a broader context, our work supports a novel pathogenic mechanism by which the nigro-striatal synapses might be affected at the early stage of the neurodegenerative process. Indeed, the LRRK2-mediated impairment of the astrocytic glutamate reuptake capacity could induce a premature striatal glutamate accumulation, anticipating subsequent irreversible consequences on the integrity of dopaminergic connections.

## Author contribution

L.I. and L.C. designed the study, optimized and performed experiments, interpreted data, and wrote the manuscript; V.G. performed imaging experiments and helped in preparing primary astrocyte cultures; F.P. and G.P. performed qPCR experiments; E.Gi. and N.P. contributed to data analysis; An.M., T.B. and G.B. performed gliosomes experiments; I.B. and G.A. performed mass spec experiments; A.D.I., C.R. and E.B. performed electrophysiological experiments; Al.M. and C.P. performed TIRFM experiments; R.B. provided human samples; E.Gi., L.B., E.Gr. and S.H. have contributed to manuscript editing and optimization. All authors have read and approved the final manuscript.

## Acknowledgments

The Authors thank the University of Padova to support L.C. as assistant professor and the IRCCS San Camillo Hospital, Venice, Italy. This work was supported by UniPD (STARs 2019: Supporting TAlents in ReSearch) and the Italian Ministry of Health (GR-2016-02363461) to L.C. L.I. is a post-doctoral fellow supported by UniPD. G.P. is supported by Fondazione Telethon (grant TDPG00514TA), MIUR (PRIN-2017ENN4FY), and Fondazione Cariplo (project 2019-3415). F.P. received support by Fondazione Caritro (project 2019.0230). R.B. is funded by Reta Lila Weston Institute and the British Neuropathological Society. S.H. is funded by the MJFF and by intramural funds from Rutgers University. We thank Dr. Alessandra Bellan for the technical support to the transfection of organotypic slices and Raffaella Cinquetti for the technical support to the electrophysiological experiments.

---

**Supplementary Figure 1.**
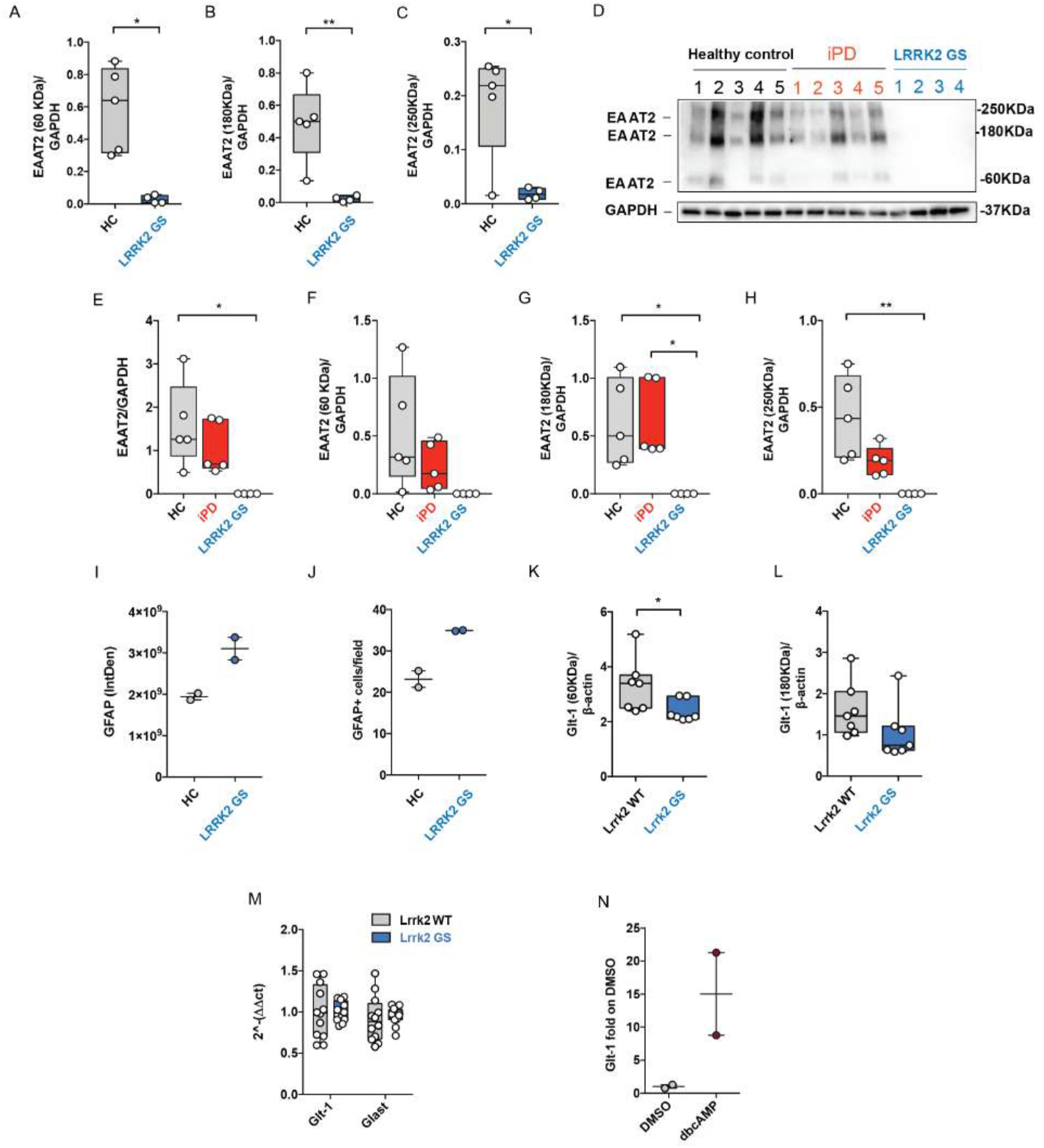
A-C) Western blot quantification of the monomeric fractions (60 KDa) as well as of the multimeric fractions (180 KDa and 250 KDa) of EAAT2 in LRRK2 G2019S PD patients (n=4) compared to healthy controls (n=5); D) Western blot analysis of healthy controls, idiopathic PD patients (iPD) and LRRK2 G2019S caudate and putamen lysates using an anti-EAAT2 antibody; E-H) Relative quantification of the total EAAT2 as well as of the monomeric fractions (60 KDa) and multimeric fractions (180 KDa and 250 KDa) of EAAT2 in the healthy controls (n=5), iPD patients (n=5) and LRRK2 G2019S PD patients (n=4). The band intensity was performed using ImageJ and normalized to the housekeeping protein GAPDH; I-J) Quantification of GFAP IntDen and of GFAP^+^ positive cells in the caudate and putamen of healthy controls (n=2) and LRRK2 G2019S PD patients (n=2); K-L) Quantification of the 60 KDa and 180 KDa Glt-1 bands in Lrrk2 G2019S and Lrrk2 WT mice; n=7 animals for each genotype; M) qPCR analysis of Glt-1 and Glast mRNA in the striatum of Lrrk2 WT and Lrrk2 G2019S mice; n=12 animals for each genotype; N) qPCR analysis of Glt-1 mRNA in primary Lrrk2 WT striatal astrocytes treated with or without dbcAMP for 10 days; experiment performed twice on n=2 independent cell cultures. Statistical analysis was performed in A-C using Unpaired T-test and in K-L using Mann-Whitney test. Statistical analysis was performed in E,G using Kruskal-Wallis test followed by Dunn’s multiple comparisons test and in F,H using One-way ANOVA test followed by Tukey’s multiple comparisons test.

**Supplementary Figure 2.**
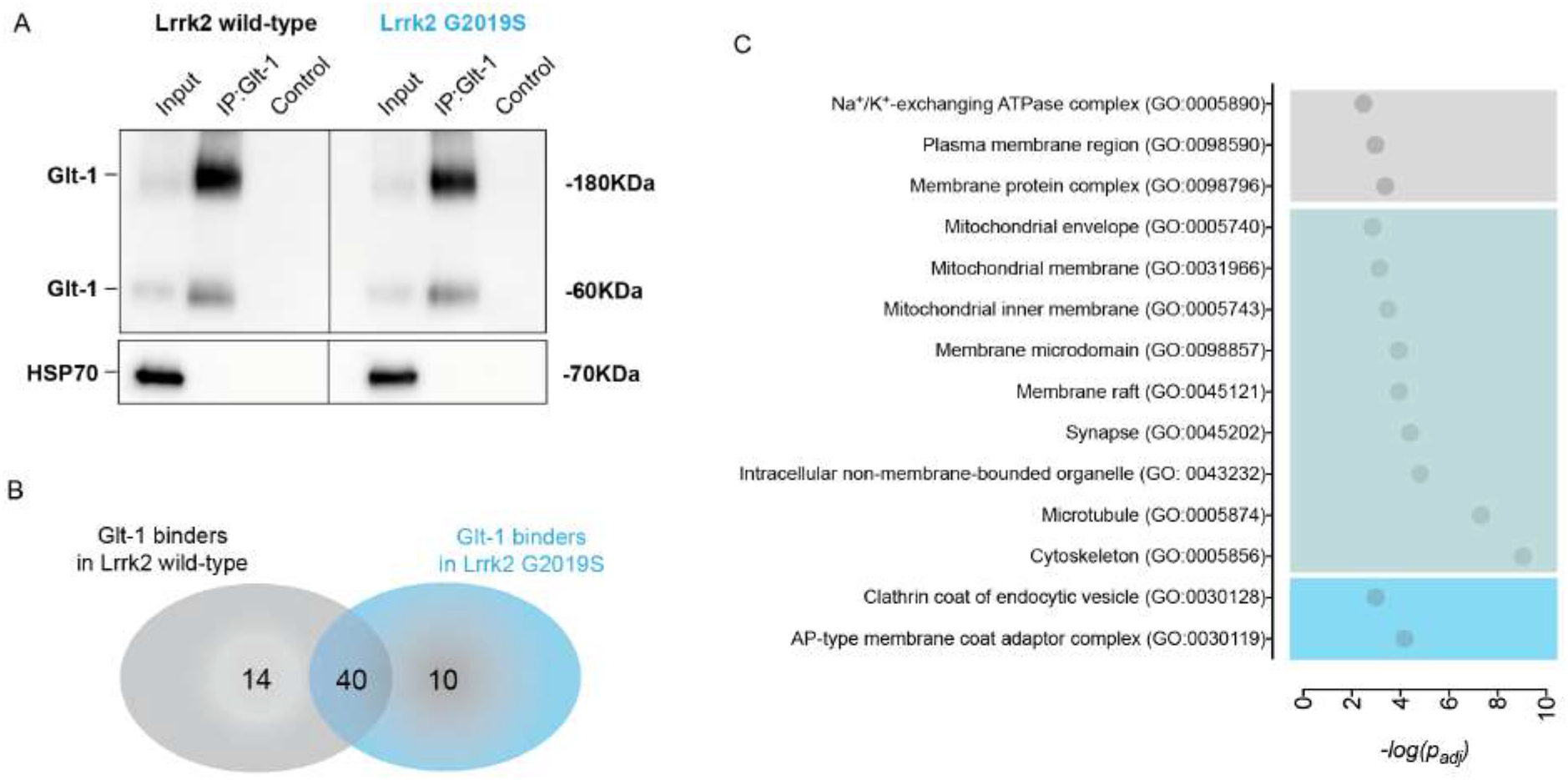
Screening of Glt-1 protein-protein interactions. A) Striatal Glt-1 immunoprecipitated from Lrrk2 WT and G2019S brains was resolved by immunoblotting. Anti-HSP70 was applied to normalize for protein content; B) Venn diagram summary of interacting proteins, colored for different background (grey for Lrrk2 WT, light blue for Lrrk2 G2019S, dark blue for Lrrk2 WT and G2019S shared interacting proteins); C) Selected GO term enrichments from CC categories using gProfiler were plotted using -log(P*adj*) values.

**Supplementary Figure 3.**
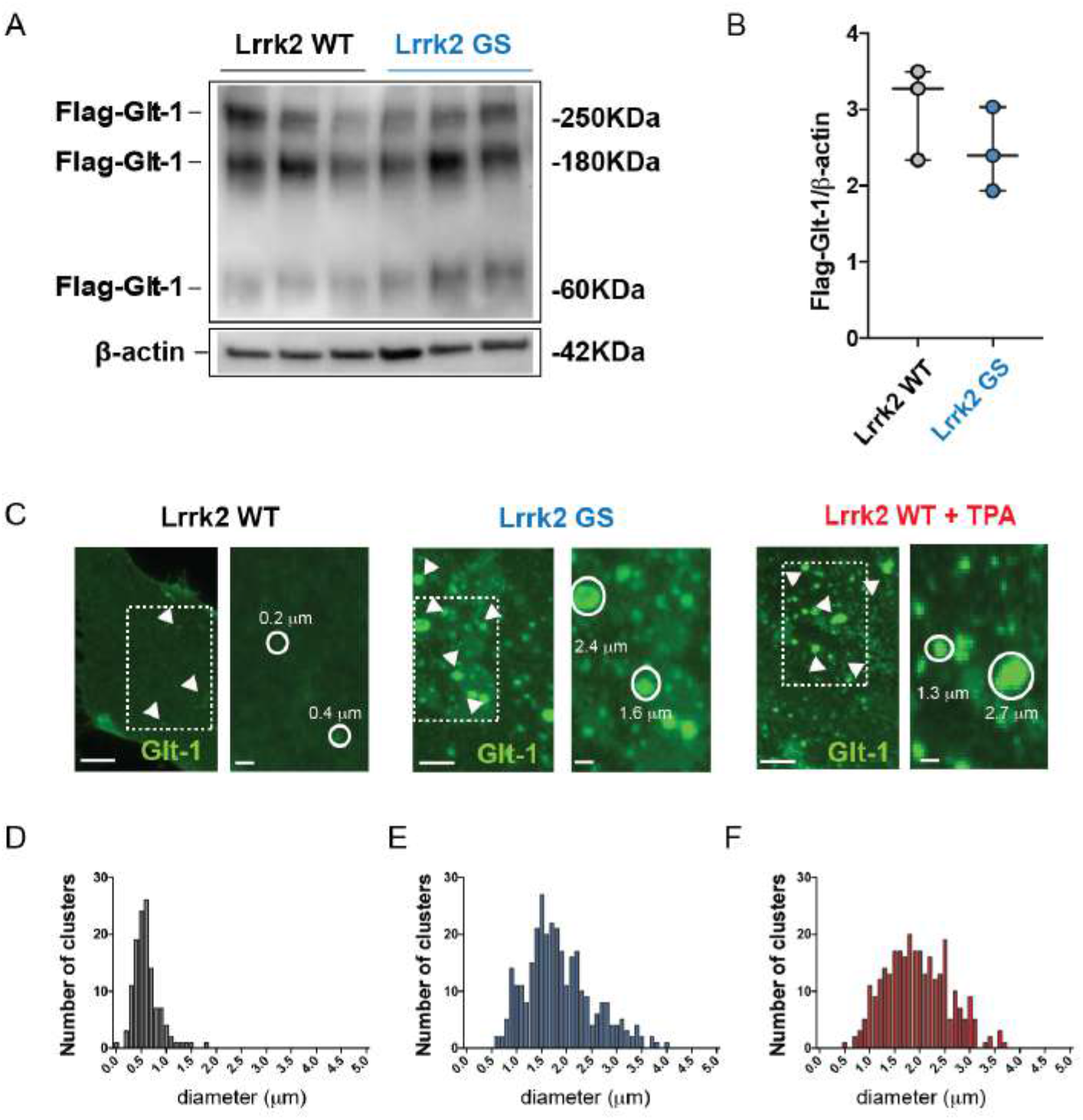
A) Western blot analysis of transfected Flag-Glt-1 expression in primary striatal Lrrk2 WT and G2019S astrocytes using an anti-Flag-HRP antibody; B) Relative quantification of band intensity was performed using ImageJ and normalized to β-actin; experiment performed in triple; C) Representative epifluorescence images of Glt-1 clusters in Flag-Glt-1 transfected primary striatal astrocytes from Lrrk2 WT (untreated or treated with TPA) and from Lrrk2 G2019S astrocytes. Scale bars 5 μm; insets 1 μm. D) Frequency distribution of Glt-1 cluster diameter in untreated Lrrk2 WT (grey bars); E) Frequency distribution of Glt-1 cluster diameter in Lrrk2 G2019S (blue bars); F) Frequency distribution of Glt-1 cluster diameter in Lrrk2 WT treated with TPA (d; red bars); n=5 cells analyzed for each group. Statistical analysis in B was performed using Mann-Whitney test.

**Supplementary Figure 4.**
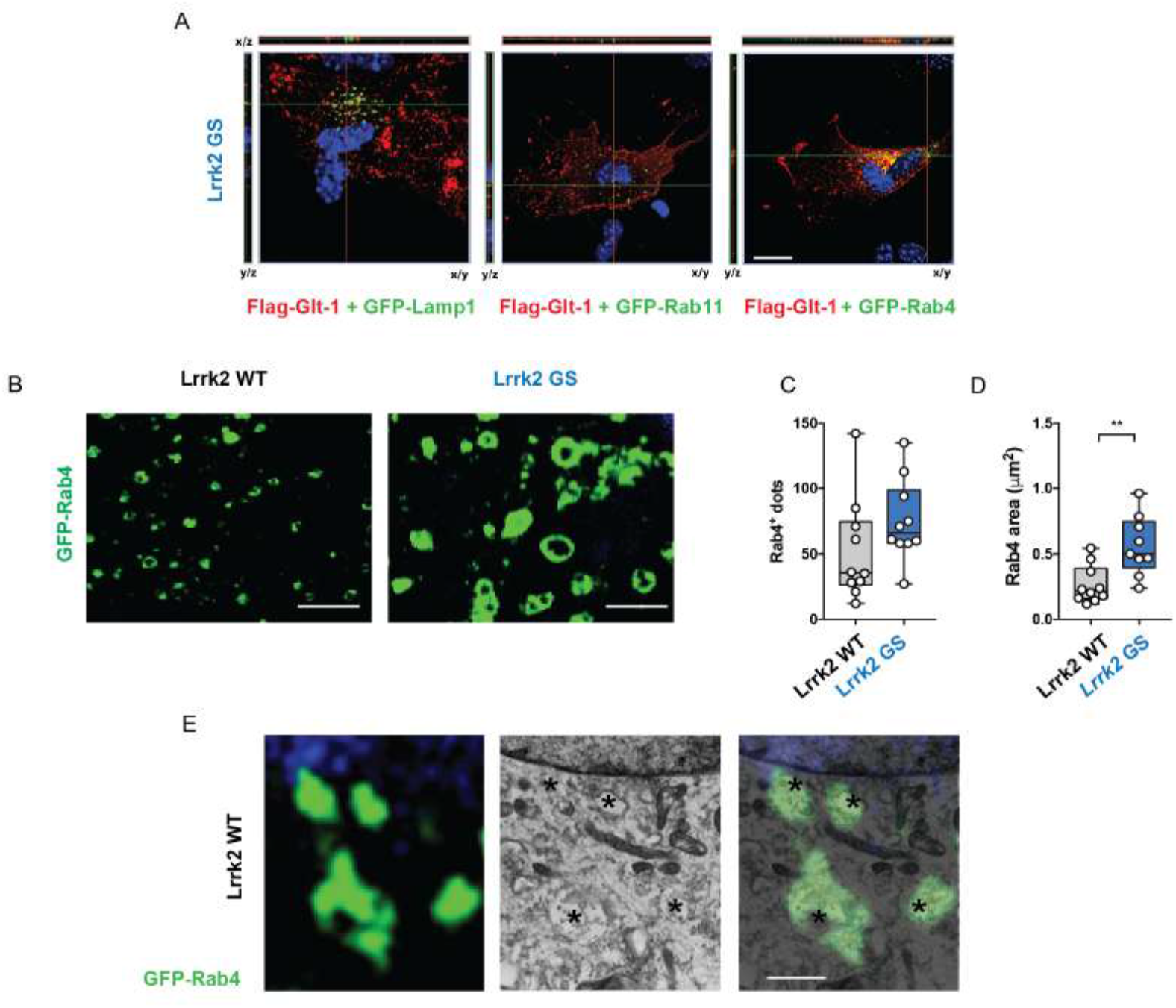
A) Orthogonal z-stack projections of Lrrk2 G2019S astrocytes co-transfected with Flag-Glt-1 (red) and GFP-Lamp1, GFP-Rab11 or GFP-Rab4 (green). Scale bar 20 μm. B) Representative confocal microscopy images of Lrrk2 WT and G2019S primary striatal astrocytes transfected with GFP-Rab4 and Flag-Glt-1. Scale bars: 2 μm; C,D) Quantification of Rab4-positive vesicle number and area; at least n=9 cells analyzed for each group and experiments performed at least in triple; E) Representative confocal (left panel), electron microscopy (middle panel) and merge (right panel) image of a Lrrk2 WT primary astrocyte transfected with Rab4-GFP (green). The image reveals the co-localization of early endosomal structures identified by transmission microscopy with the GFP-Rab4-positive vesicles identified by confocal microscopy; scale bar: 1 μm. Statistical analysis in C,D was performed using Unpaired t-tests.

**Supplementary Figure 5.**
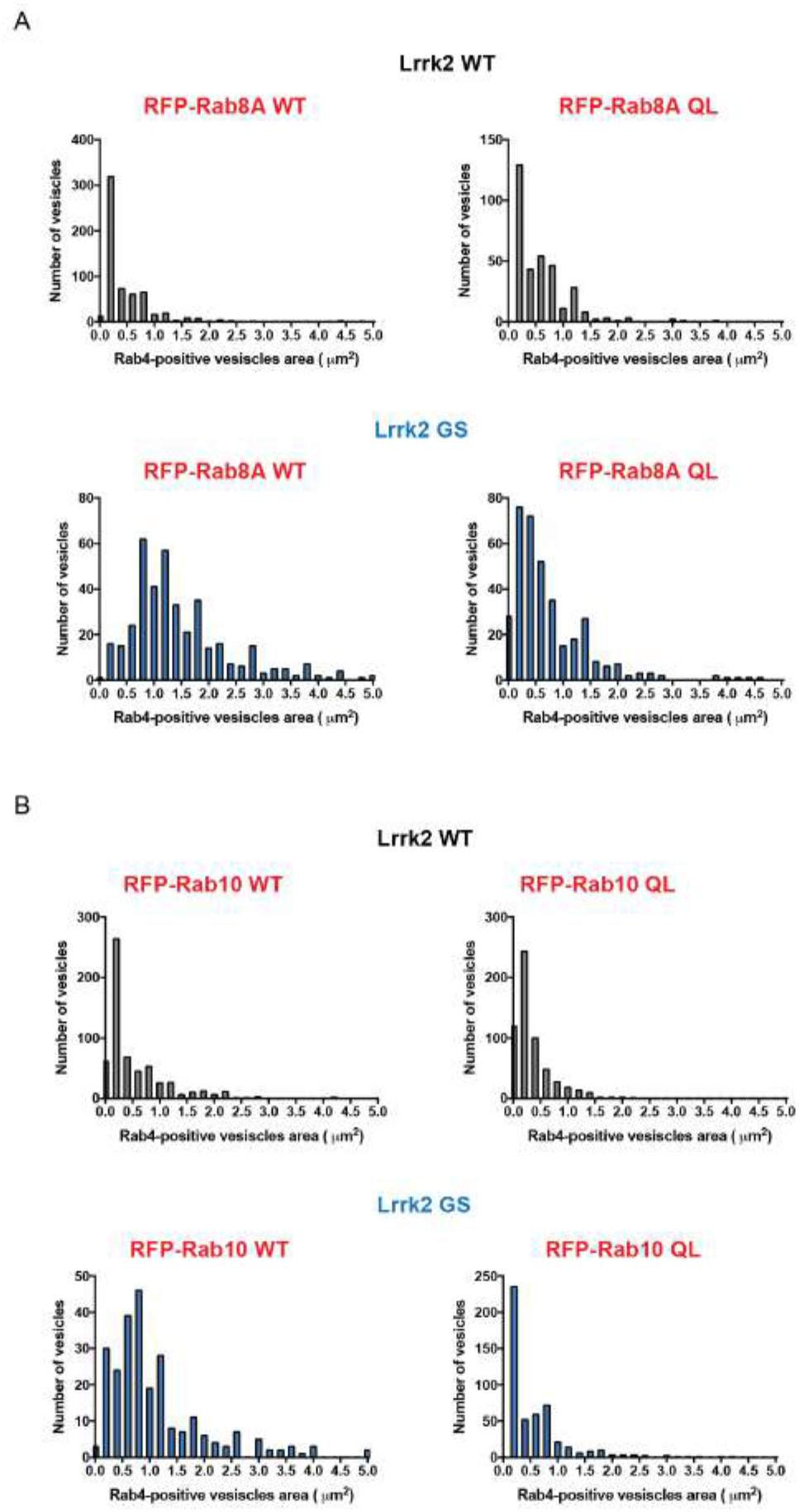
A,B) Frequency distribution of Rab4-positive vesicles area in Lrrk2 WT and Lrrk2 G2019S astrocytes. At least n=6 cells analyzed for each group and experiments performed at least in triple.

**Supplementary Figure 6.**
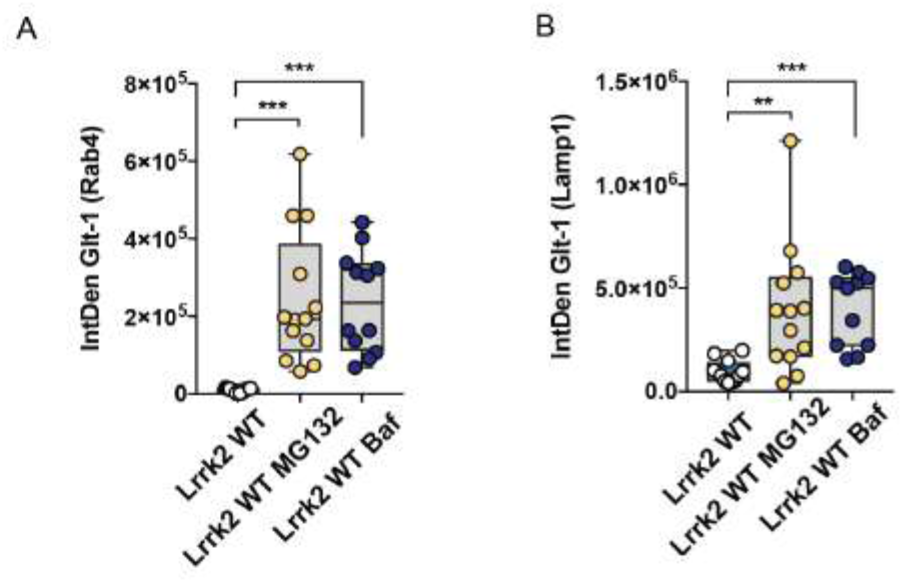
A) Quantification of Glt-1 IntDen in the Rab4-positive vesicles in Lrrk2 WT astrocytes under basal conditions (n=13 cells) or upon MG132 (n=13 cells) or Bafilomycin application (n=12 cells); B) Quantification of Glt-1 IntDen in the endogenous Lamp1-positive structures in Lrrk2 WT astrocytes under basal conditions (n=12 cells) or upon MG132 (n=13 cells) or Bafilomycin (n=11 cells) application; experiments performed at least in triple. Statistical analysis in A was performed using One-way ANOVA test followed by Tukey’s multiple comparisons test. Statistical analysis in B was performed using Kruskal-Wallis test followed by Dunn’s multiple comparisons test.

**Supplementary File 1**

**Supplementary File 2**

**Supplementary File 3**

**Supplementary File 4**

